# Self-organizing ovarian somatic organoids preserve cellular heterogeneity and reveal cellular contributions to ovarian aging

**DOI:** 10.1101/2024.08.10.607456

**Authors:** Shweta S. Dipali, Madison Q. Gowett, Pratik Kamat, Aubrey Converse, Emily J. Zaniker, Abigail Fennell, Teresa Chou, Michele T. Pritchard, Mary Zelinski, Jude M. Phillip, Francesca E. Duncan

## Abstract

Ovarian somatic cells are essential for reproductive function, but no existing *ex vivo* models recapitulate the cellular heterogeneity or interactions within this compartment. We engineered a novel ovarian somatic organoid model by culturing a stroma-enriched fraction of mouse ovaries in scaffold-free agarose micromolds. Ovarian somatic organoids self-organized, maintained diverse cell populations, produced extracellular matrix, and secreted hormones. Organoids generated from reproductively old mice exhibited reduced aggregation and growth compared to young counterparts, as well as differences in cellular composition. Interestingly, matrix fibroblasts from old mice demonstrated upregulation of pathways associated with the actin cytoskeleton and downregulation of cell adhesion pathways, indicative of increased cellular stiffness which may impair organoid aggregation. Cellular morphology, which is regulated by the cytoskeleton, significantly changed with age and in response to actin depolymerization. Moreover, actin depolymerization rescued age-associated organoid aggregation deficiency. Overall, ovarian somatic organoids have advanced fundamental knowledge of cellular contributions to ovarian aging.

## Introduction

Ovarian follicles, comprised of oocytes and their surrounding somatic support cells, are the functional units of the ovary and are essential for producing gonadal hormones as well as fertilization-competent gametes. Ovarian follicles reside and develop in a complex and heterogenous stromal microenvironment composed of fibroblasts, immune cells, epithelial cells, interstitial fibroblasts and mesenchymal cells, blood and lymphatic vessels, nerves, and a dynamic extracellular matrix (ECM) ^1^. The stromal microenvironment is critical in supporting follicle development as co-culture of primary and early secondary follicles with feeder layers of macrophage and theca-enriched stromal cells improves folliculogenesis endpoints ^2^.

The structure and function of the ovarian stroma undergoes significant alterations in various conditions, including polycystic ovary syndrome, ovarian cancer, and reproductive aging ^1, 3, 4^. Ovaries from reproductively old mice have a greater proportion of immune and epithelial cells, as well as fewer broadly classified fibroblastic stromal cells than young counterparts ^5^. Additionally, the ovarian stroma becomes inflammatory with advanced age and is associated with altered macrophage ontogeny and polarization, as well as the presence of a unique population of multinucleated macrophage giant cells ^6–8^. There is also an age-dependent shift in ovarian immune cell populations with a skew towards adaptive immunity ^5, 9^. The relative abundance of subpopulations of ovarian fibroblasts changes with age, as does their transcriptome ^5, 10^. Reproductive aging results in greater proportions of senescence-associated secretory phenotype (SASP) fibroblasts in the ovarian stroma, as well as decreased expression of genes involved in collagen degradation, which may contribute to age-related ovarian fibrosis and tissue stiffness ^5, 6, 10–12^. In addition to collagens, the composition of the broader ovarian ECM is altered with age ^13, 14^.

Given the importance of the stroma in ovarian physiology, pathology, and aging, robust *in vitro* strategies are needed to study this compartment in a controlled manner. However, unlike the numerous methods that exist to culture ovarian follicles, there are few *in vitro* models of the ovarian somatic compartment, and those that exist have limitations in capturing the diversity of cell types and interactions present ^15^. Several methods exist to culture specific ovarian cell populations, such as primary mouse ovarian surface epithelial cells, but these models inherently do not capture the cellular heterogeneity and diverse cell-cell and cell-matrix interactions of the ovarian stroma *in vivo* ^16^. Additionally, although primary somatic cells isolated from a stroma-enriched fraction of the mouse ovary are initially heterogenous, two-dimensional culture using traditional methods results in expansion of the macrophage population ^2^. Ovarian tissue explants maintain the cellular heterogeneity and structural organization of the native tissue and have been used to investigate ovarian physiology and pathologies, but they can vary in composition due to where within the tissue the explants were derived ^17, 18^. Organoids, defined as three-dimensional, multi-cellular, miniaturized versions of organs or tissues, have been utilized to model several reproductive tissues and address the shortcomings of traditional cell culture and tissue explants ^19, 20^. However, current ovarian organoid models utilize a single cell type, model follicle structures rather than the ovarian stroma, or are focused on disease states such as ovarian cancer ^21–23^.

Therefore, the goals of this study were to engineer ovarian somatic organoids in a manner that preserves the cellular composition and function of the native tissue and to utilize this organoid model to inform mechanisms of ovarian aging. Using scaffold-free agarose micromolds, we demonstrated that a stroma-enriched fraction of ovarian somatic cells can self-assemble into solid structures. Organoids contained all major somatic cell types of the ovary throughout culture, produced an ECM, and secreted hormones. However, ovarian somatic organoids generated using cells from reproductively old mice exhibited compromised aggregation, growth, and hormone production relative to young counterparts. Such age-dependent differences in aggregation capacity were in part due to changes at the cellular level because, compared to young mice, stroma/mesenchyme cells from reproductively old mice showed an upregulation of genes enriched in actin-related pathways and downregulation of genes enriched in cell adhesion and migration pathways. Together, these findings are suggestive of an age-dependent increase in cellular stiffness which may contribute to decreased organoid aggregation. The actin cytoskeleton also plays an important role in regulating cell morphology. Analysis at single cell resolution using a machine-learning methodology revealed that primary ovarian somatic cells have heterogenous morphology that changes with age. Depolymerization of the actin cytoskeleton using Latrunculin A shifted ovarian somatic cell morphology and restored the aggregation potential of organoids from old mice. Thus, this robust organoid model may be used to study the ovarian somatic compartment and has revealed novel cellular mechanisms underlying aging in the ovarian stroma. Importantly, we can also generate ovarian somatic organoids with rhesus macaque ovarian tissue, enabling future translation to primate models.

## Results

### Ovarian somatic organoids self-assemble and contain key ovarian cell populations with distinct spatial organization

Mouse ovarian somatic organoids were generated by digesting an enriched ovarian stromal fraction into a single cell suspension followed by filtering and differential plating to isolate viable primary ovarian somatic cells from oocytes (Figure 1A). These cells were then seeded into agarose micromolds to generate organoids in an ECM scaffold-free manner (Figure 1A). IntestiCult mouse intestinal organoid growth medium best supported organoid aggregation and growth relative to other commercially available media, including HepatiCult, MesenCult, and RPMI-1640 (Figure 1B-D). The resulting organoids were solid structures that remained intact and continued to grow even when removed from the agarose micromolds (Figure 1B, Supplemental Figure S1A-B). Furthermore, organoids exhibited similar percentages of proliferating and apoptotic cells as the mouse ovary *in vivo* (organoids Ki67-positive area: 15.7 ± 0.8%, ovary Ki67-positive area: 9.5 ± 2.1%, P>0.05; organoids CC3-positive area: 2.6 ± 0.2%, ovary CC3-positive area: 1.6 ± 0.6%, P>0.05) (Figure 1E-G). Interestingly, while Ki67-positive cells were distributed throughout the organoids, CC3-positive cells were primarily restricted to the perimeter of organoids (Figure 1E, I). Of note, these methods were also used to generate organoids using primary ovarian somatic cells isolated from both the cortex and medulla of rhesus macaque ovaries (Supplemental Figure S1C-D). Thus, this novel technique has translatability to non-human primates, humans, and other large mammalian species To determine whether key ovarian cell types were present in these organoids, we performed immunohistochemistry using antibodies against vimentin (fibroblasts and myofibroblasts), alpha-smooth muscle actin (11-SMA; myofibroblasts and smooth muscle cells), desmin (smooth muscle cells), F4/80 (murine macrophages), FOXL2 (granulosa and granulosa-lutein cells), and 3-beta-hydroxysteroid dehydrogenase (3β-HSD; theca, theca-lutein, and steroidogenic stromal cells) (Figure 1H; Supplemental Figure S2). By day 6 of culture, vimentin-positive fibroblasts and FOXL2-positive steroidogenic cells were present throughout ovarian somatic organoids, whereas myofibroblasts and smooth muscle cells marked by 11-SMA and desmin were not present in the organoids (Figure 1H-I; Supplemental Figure S2A-C, E). F4/80-positive macrophages were restricted to the organoid perimeter similar to CC3-positive apoptotic cells (Figure 1H-I; Supplemental Figure S2D). Interestingly, the F4/80-positive cells were hypertrophied by day 6 of culture consistent with phagocytic activity (Figure 1H-I; Supplemental Figure S2D). Moreover, 3β-HSD-positive cells self-organized such that they were excluded from the organoid core (Figure 1H-I; Supplemental Figure S2F).

**Figure 1.**
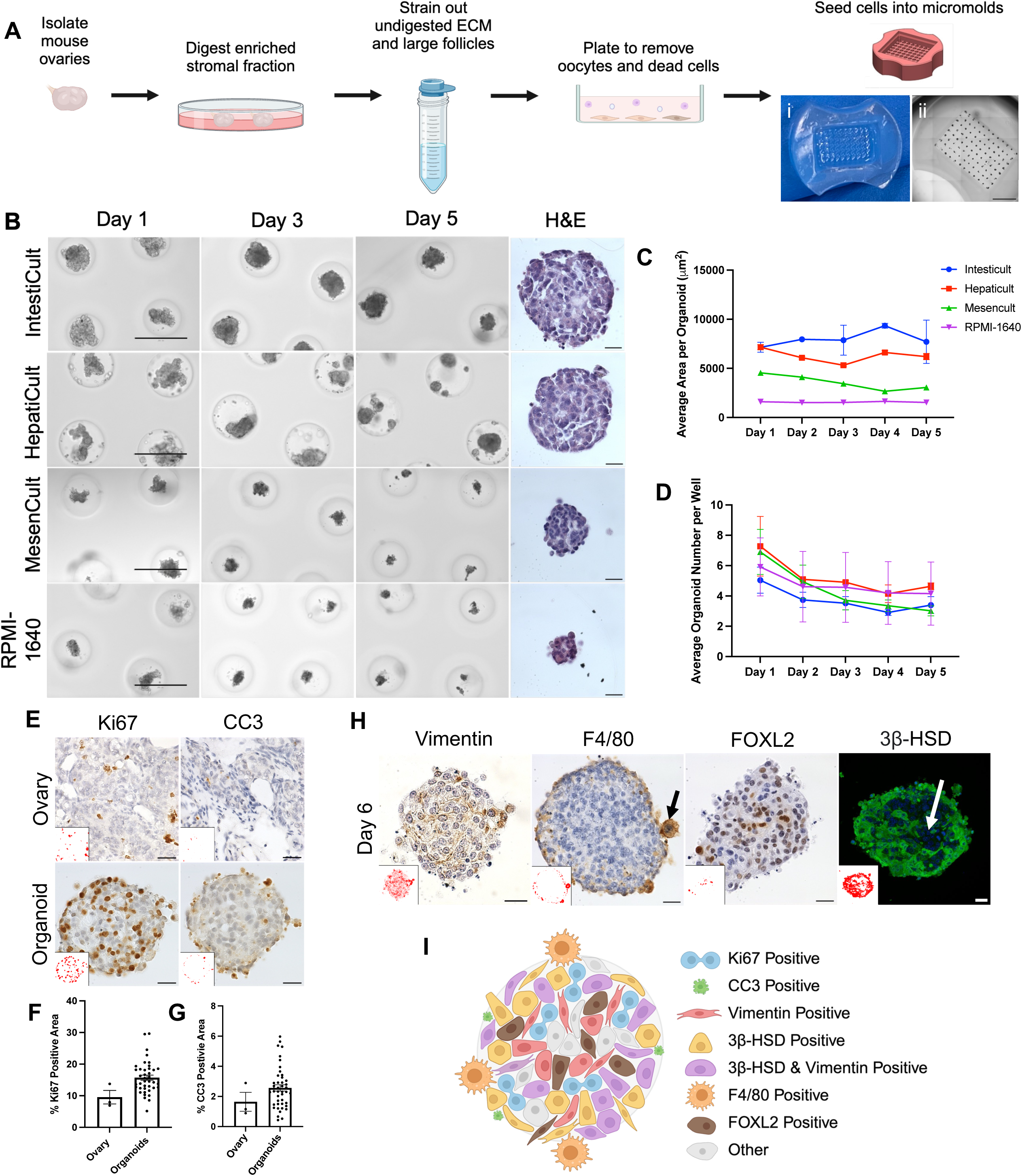
Ovarian somatic organoids self-assemble into solid structures containing key ovarian cell populations with distinct spatial organization. A) Schematic of workflow to isolate primary mouse ovarian somatic cells and generate organoids and images of an empty agarose micromold (i) and a scan of an agarose micromold following organoid generation (ii). B) Representative images of ovarian somatic organoids cultured in IntestiCult, HepatiCult, MesenCult, or RPMI-1640 media at days 1, 3, and 5 of culture. Representative images of H&E-stained organoid sections following 5 days in culture in each media. Scale bars for transmitted light images = 400 µm and scale bars for H&E images = 20 µm. C-D) Quantification of the average area per organoid (C) and average number of organoids per microwell (D) over 5 days of culture in each media. N=3-4 micromolds per media type. E) Representative images of Ki67 and cleaved caspase-3 (CC3) IHC staining of mouse ovarian tissue sections and ovarian somatic organoid sections. Scale bars = 20 µm. Insets are thresholded images that show positive staining. F-G) Quantification of the percent Ki67 (F) and CC3 (G) positive area in regions of interest in mouse ovaries (N=3 ovaries) and ovarian somatic organoids after 5 days of culture (N=40-44). H) Representative images of IHC labelling of organoid sections after 6 days of culture using antibodies against Vimentin, F4/80, FOXL2, and 3β-HSD. Arrows show F4/80-positive macrophages restricted to the perimeter of organoids (black) and 3β-HSD-positive steroidogenic cells absent from the core of organoids (white). Insets are thresholded images that show positive staining. Scale bars = 20 µm. I) Schematic depicting the presence and spatial organization proliferative (Ki67-positive) and apoptotic (CC3-positive) cells, as well as key ovarian cell populations within organoids.

Significantly fewer vimentin-positive fibroblasts were present in organoids at both days 1 (23.8 ± 1.3%, P=0.04) and 6 (24.6 ± 2.1%, P=0.004) of culture compared to the mouse ovary (48.7 ± 10.3%) (Supplemental Figure S2A). Although not statistically significant, fewer 11-SMA-positive cells were present in organoids at day 1 of culture (5.5 ± 0.8%) compared to the mouse ovary (14.2 ± 2.3%), and these cells were mostly absent in organoids by day 6 of culture (0.6±0.2%, P<0.0001) in comparison to day 1 (Supplemental Figure S2B). Additionally, desmin-positive cells were largely absent in organoids at days 1 (0.2 ± 0.1%) and 6 (0.1 ± 0.0%) of culture (Supplemental Figure S2C). However, by day 6 of culture, F4/80-postive macrophages, in addition to FOXL2- and 3β-HSD-positive steroidogenic cells were present in organoids in similar quantities to the mouse ovarian somatic compartment *in vivo* (Supplemental Figure S2D-F). Overall, ovarian somatic organoids are composed of diverse cell populations which undergo dynamic changes during culture.

### Organoids exhibit age-dependent differences in aggregation, growth, and function

Given that the ovarian stroma changes significantly with age *in vivo*, we wanted to determine whether age-dependent differences were reflected in the ability of primary ovarian somatic cells to form organoids ^5, 13^. Organoids generated from reproductively old mice exhibited impaired aggregation as determined by quantifying the average number of aggregates per microwell, with a higher number indicating poor aggregation potential (Figure 2A-B). There were more aggregates per microwell of the agarose molds in samples derived from reproductively old mice relative to young at both day 1 and 2 of culture (old day 1: 6.2 ± 0.6, young day 1: 2.2 ± 0.5, P<0.0001; old day 2: 4.6 ± 0.6, young day 2: 1.7 ± 0.2, P<0.0001) (Figure 2A-B). Additionally, organoids generated from old mice were 46.6 - 61.7% smaller in size across six days of culture compared to organoids from young mice (Figure 2A, C). Consistent with this smaller size, organoids generated from old mice had fewer Ki67-positive proliferative cells and more CC3-positive apoptotic cells than young counterparts (old Ki67: 4.6 ± 0.7%, young Ki67: 7.4 ± 1.0%, P=0.03; old CC3: 1.5 ± 0.2%, young CC3: 0.4 ± 0.03%, P<0.0001) (Figure 2D-E).

**Figure 2.**
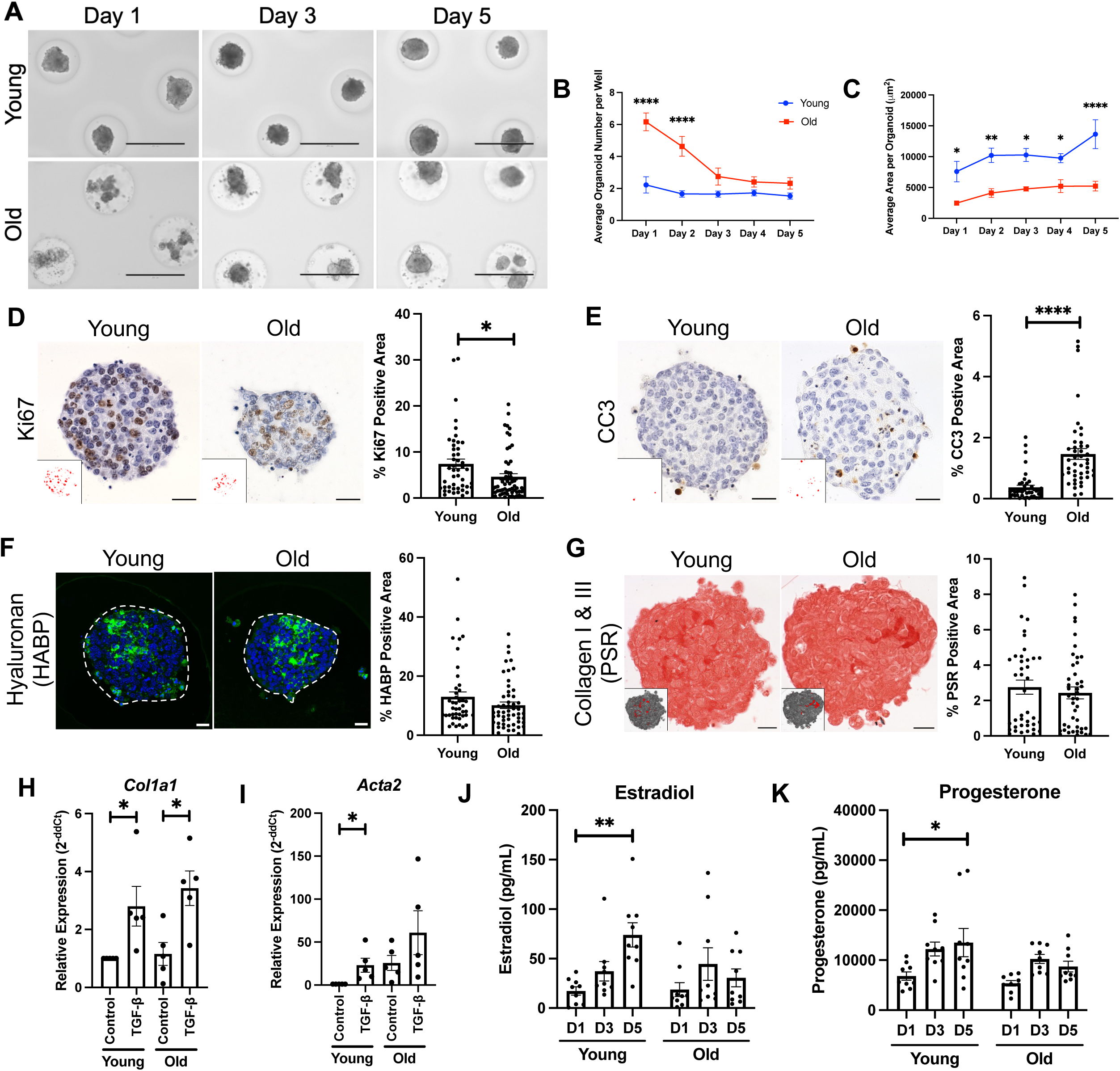
Organoids exhibit age-dependent differences in aggregation, growth, and function. A) Representative images of ovarian somatic organoids generated from reproductively young (6-12 weeks) or reproductively old (10-14 months) mice at days 1, 3, and 5 of culture. Scale bars = 400 µm. B-C) Quantification of the average number of organoids per microwell (B) and average area per organoid (C) over 5 days of culture for organoids generated from reproductively young and old mice. *P<0.05, **P<0.01, and ****P<0.0001 by two-way ANOVA. N=4 micromolds per age. D-E) Representative images of Ki67 (D) and cleaved caspase-3 (CC3, E) IHC staining of young and old ovarian somatic organoid sections. Scale bars = 20 µm. Insets are thresholded images that show positive staining. Quantification of the percent Ki67 (D) and CC3 (E) positive area in young (N=42-46) and old (N=44-57) ovarian somatic organoids after 5 days of culture. *P<0.05 and ****P<0.0001 by Welch’s t-tests. F) Representative images of hyaluronan binding protein (HABP, green) assays performed with young and old ovarian somatic organoid sections. Nuclei (blue) were detected with DAPI. Scale bars = 20 µm. Organoids are outlined by dashed lines. Quantification of the percent HABP-positive area in young (N=43) and old (N=53) ovarian somatic organoids after 5 days of culture. G) Representative images of Picrosirius Red (PSR)-stained young and old ovarian somatic organoid sections. Scale bars = 20 µm. Insets are thresholded images that show positive staining. Quantification of the percent PSR-positive area in young (N=43) and old (N=53) ovarian somatic organoids after 5 days of culture. H-I) Gene expression of *Col1a1* (H) and *Acta2* (I) in ovarian somatic organoids measured by RT-qPCR following 48-hour treatment with 10 ng/mL TGF-β or control. Gene expression for TGF-β-treated organoids was graphed as fold-change over young. *P<0.05 by t-tests. N=5 trials. J-K) Estradiol (J) and progesterone (K) secretion measured in conditioned media of young and old ovarian somatic organoids by ELISAs at days 1, 3, and 5 of culture. *P<0.05 and **P<0.01 by two-way ANOVAs. N=9 micromolds over 3 trials.

The extracellular matrix (ECM) is a critical component of the ovarian stroma, with hyaluronan and collagen being two major constituents ^1, 11, 24^. Using a hyaluronan binding protein assay and Picrosirius Red staining, we demonstrated that organoids produce both hyaluronan (young: 13.0 ± 1.7%, old: 10.2 ± 1.1%, P>0.05) and collagen I & III (young: 2.8 ± 0.4%, old: 2.4 ± 0.3%, P>0.05) (Figure 2F-G). Of note, organoids trended towards the age-dependent decrease in hyaluronan that occurs in mouse and human ovaries, but did not exhibit the characteristic age-associated increase in collagen (Figure 2F-G) ^6, 10–12^. To determine, however, if the organoids were responsive to a pro-fibrotic stimulus, we treated them with transforming growth factor-beta (TGF-β), which has been demonstrated to induce collagen expression in several cell types including cardiac fibroblasts, alveolar fibroblasts, and myoblasts, as well as primary ovarian somatic cells cultured in a two-dimensional monolayer ^25, 26^. Following TGF-β treatment we assessed expression of *Col1a1* and *Acta2*, which encode one chain of collagen I and 11-SMA, respectively, and are upregulated in fibrotic processes (Figure 2H-I) ^27, 28^.

TGF-β treatment increased expression of *Col1a1* and *Acta2* in organoids generated from reproductively young (*Col1a1*: 2.8 ± 1.5-fold increase, P=0.03, *Acta2*: 23.1 ± 18.3-fold increase, P=0.03) and old (*Col1a1*: 3.0 ± 0.5-fold increase, P=0.01, *Acta2*: 2.4 ± 1.0-fold increase, P>0.05) mice, demonstrating the functional responsiveness of these organoids (Figure 2H-I).

Given that steroid hormone production is an important role of the ovary, we evaluated estradiol and progesterone production by ovarian somatic organoids at days 1, 3, and 5 of culture to further probe their functional capacity (Figure 2J-K). Organoids generated from both young and old mice produced estradiol and progesterone.

However, while organoids from young mice demonstrated a significant increase in estradiol (Day 1: 17.3 pg/mL, Day 5: 73.9 pg/mL, P=0.005) and progesterone (Day: 6813.4 pg/mL, Day 5: 13507.9 pg/mL, P=0.03) secretion over the culture period, organoids generated from old mice did not, indicative of impaired endocrine output as expected with age (Figure 2J-K) ^29^.

### Heterogeneous cell populations are maintained in ovarian somatic organoids irrespective of age

To obtain a more comprehensive understanding of the cell populations present within these ovarian somatic organoids throughout culture and determine whether any age-dependent differences exist, we utilized single cell transcriptomic analysis. Single cell RNA-sequencing (scRNAseq) was performed using primary ovarian somatic cells isolated from both reproductively young and old mice following plating but prior to organoid generation (monolayer), as well as from organoids generated from young and old mice on day 1 and day 6 of culture (Figure 3A). Unbiased clustering and uniform manifold approximation and projection (UMAP) analysis revealed 11 bioinformatically distinct cell clusters that were subsequently identified manually using known markers of mouse ovarian cell types (Figure 3B, Supplemental Figure S3A-B) ^5, 30, 31^. Luteal cells bioinformatically segregated into two clusters and were the most abundant cell type present (*n*=27,269 cells) followed by stroma/mesenchyme cells (*n*=15,235), and epithelial cells (*n*=10,923) (Figure 3B). Other cell clusters included granulosa, theca, immune, endothelial, and mitotic cells (Figure 3B). The relative abundance of these cell types was as expected given our use of a stroma-enriched fraction of the ovary.

**Figure 3.**
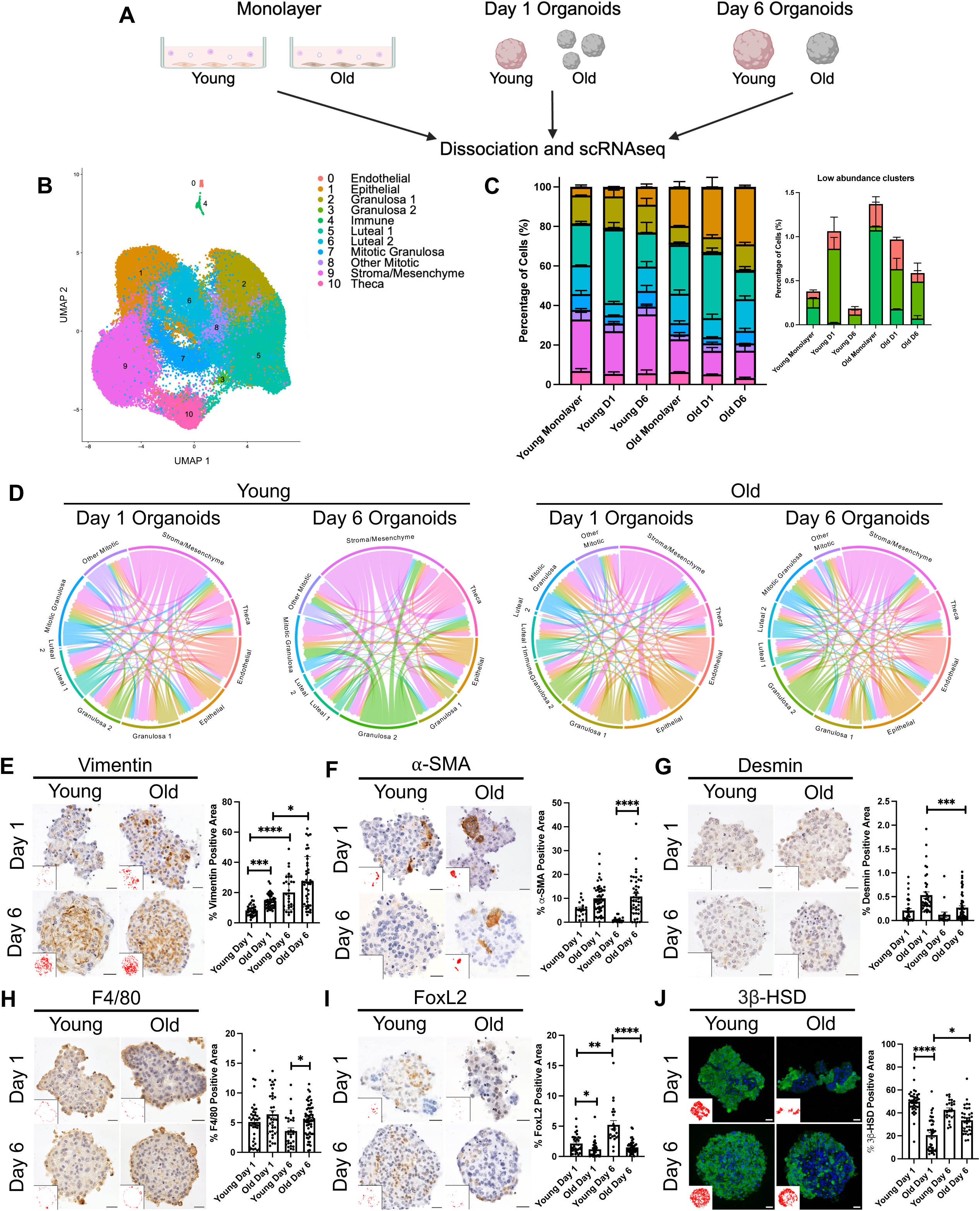
Heterogenous cell populations are maintained in ovarian somatic organoids irrespective of age, over six days in culture. A) Schematic of samples utilized for single cell RNA-sequencing (scRNAseq). B) Uniform manifold approximation and projection (UMAP) plot of primary mouse ovarian somatic cells from reproductively young (6-12 weeks) and reproductively old (10-14 months) mice following plating in 2D, prior to organoid generation (monolayer) and cells dissociated from young and old ovarian somatic organoids at days 1 and 6 of culture combined. Unbiased clustering revealed 11 distinct cell populations. scRNAseq was performed for 2 replicates per group. C) Quantification of the percentage of cells in each cluster for monolayer, day 1 organoids, and day 6 organoids for each age. Inset shows percentage of cells in low abundance clusters. Cluster color coding as in B. D) Chord diagrams showing cell-cell communication in day 1 and day 6 organoids for each age created using CellChat. The color of the chords indicate the cell cluster sending the signal and the size of the chords are proportional to signal strength. E-J) Representative images of vimentin (E), alpha-smooth muscle actin (11-SMA, F), desmin (G), F4/80 (H), FOXL2 (I), and 3β-HSD (J) IHC staining of young and old ovarian somatic organoid sections at days 1 and 6 of culture. Scale bars = 20 µm. Insets are thresholded images that show positive staining. Quantification of the positive area for each marker in young and old ovarian somatic organoids at days 1 and 6 of culture (Young Day 1: N=16-41, Old Day 1: N=34-50, Young Day 6: N=15-29, Old Day 6: N=37-59). *P<0.05, **P<0.01, ***P<0.001, and ****P<0.0001 by Kruskal-Wallis tests followed by Dunn’s multiple comparisons tests. Statistical significance only for Young Day 1 vs Old Day 1, Young Day 6 vs Old Day 6, Young Day 1 vs Young Day 6, and Old Day 1 vs Old Day 6 is shown on graphs. All other statistically significant comparisons are listed in Supplemental Table S1.

Importantly, these cell clusters were present in all organoids irrespective of age group and time in culture, demonstrating the ability of our organoid model to maintain cellular heterogeneity *in vitro* (Figure 3C, Supplemental Figure S3A, C). Nevertheless, the relative abundance of certain cell clusters shifted with age (Figure 3C, Supplemental Figure S3A, C). For example, relative to young counterparts, primary ovarian somatic cells and organoids derived from reproductively old mice had a 3.2–5.5-fold greater percentage of epithelial cells and a 4.5-6.4-fold greater percentage of immune cells (Figure 3C, Supplemental Figure S3A, C). However, immune cells in general were present in low numbers (Figure 3C, Supplemental Figure S3A, C). Primary cells and organoids from old mice also contained smaller relative proportions of granulosa 1 cells (0.5–0.9-fold change) and stroma/mesenchyme cells (0.5–0.6-fold change) than those from young mice (Figure 3C, Supplemental Figure S3A, C).

To determine which cell clusters exhibited the greatest age-dependent changes in gene expression, we graphed the total number of differentially expressed genes (DEGs) with age per cluster for cells dissociated from organoids at day 6 of culture (Supplemental Figure S3D). Epithelial (*n*=959 DEGs), luteal 2 (*n*=863 DEGs), and granulosa 1 (*n*=597 DEGs) cells demonstrated the most significant age-associated changes in gene expression (Supplemental Figure S3D). Gene Ontology analysis of genes differentially expressed in the epithelial cluster with age revealed downregulation of pathways associated with cell proliferation and bone-related processes (Supplemental Figure S3E). Pathways involved in response to oxidative stress and mitosis were downregulated with age in luteal 2 cells (Supplemental Figure S3E).

Pathways downregulated in granulosa 1 cells with age included those related to cell adhesion, cell migration, inflammatory response, as well as neuron-associated pathways, which are commonly enriched in cumulus cells (Supplemental Figure S3E) ^32^.

Pathways related to aortic valve development, driven by Wnt-signaling genes, were upregulated in granulosa 1 cells with age and no pathways were significantly upregulated in epithelial or luteal 2 cells with age (Supplemental Figure S3E).

CellChat analysis revealed cell communication and integration among the various cell types in ovarian somatic organoids. Consistent with previous studies of the aging mouse ovary *in vivo*, organoids from old mice had a greater number intercellular interactions than young counterparts at both day 1 (old: 3769 interactions, young: 3511 interactions) and day 6 (old: 4477 interactions, young: 3233 interactions) of culture (Figure 3D) ^5^. At day 1 of culture, the stroma/mesenchyme cluster had the greatest number of interactions in organoids derived from young mice, whereas the epithelial cluster had the greatest number of interactions in those derived from old mice (Figure 3D). However, stroma/mesenchyme cells were the most active in intercellular signaling in both organoids derived from reproductively young and old mice by day 6 of culture (Figure 3D). Overall, cell-cell communication within organoids suggests that in addition to maintaining cellular heterogeneity, this culture method maintains cellular signaling capacity.

To further investigate age-dependent differences in cell composition and organization of ovarian somatic organoids at the protein level, we performed immunohistochemistry using antibodies against the same markers of ovarian cell types as in Figure 1. Organoids contained fibroblasts, myofibroblasts, macrophages, and steroidogenic cells irrespective of age, but had age-associated differences in the relative proportions of some cell populations (Figure 3E-J). Organoids generated from old mice exhibited a greater proportion of vimentin-positive fibroblast-lineage cells at day 1 but not at day 6 of culture compared to young counterparts (old day 1: 14.3 ± 0.7%, young day 1: 8.2 ± 0.6%, P=0.0002; old day 6: 27.4 ± 2.5%, young day 6: 20.1 ± 2.1%, P>0.05) (Figure 3E). Moreover, organoids from old mice contained significantly more 11-SMA-positive cells and F4/80-positive macrophages than young counterparts at day 6 of culture (old 11-SMA: 10.8 ± 1.2%, young 11-SMA: 1.1 ± 0.3%, P<0.00001; old F4/80: 5.5 ± 0.3%, young F4/80: 3.6 ± 0.5%, P=0.02) (Figure 3F, H). Although 11-SMA marks both myofibroblasts and smooth muscle cells, organoids derived from both young and old mice at both days 1 and 6 of culture only contained between 0.1 ± 0.1–0.5 ± 0.1% of desmin-positive smooth muscle cells, suggesting that 11-SMA-positive cells in ovarian somatic organoids are largely myofibroblasts (Figure 3G). FOXL2-positive granulosa and granulosa-lutein cells were less abundant in organoids from old mice at day 1 and day 6 of culture compared to those from young mice (old day 1: 1.2 ± 0.2%, young day 1: 2.1 ± 0.3%, P=0.02; old day 6: 1.5 ± 0.2%, young day 6: 5.2 ± 0.7%, P<0.0001) (Figure 3I). Similar to FOXL2-positive cells, 3β-HSD-positive steroidogenic cells were also less abundant in organoids from old mice at day 1, in comparison to organoids generated from young mice at day 1, but were present in organoids from young and old mice in similar quantities by day 6 (old day 1: 20.6 ± 2.2%, young day 1: 49.7 ± 1.7%, P<0.0001; young day 6: 42.6 ± 1.8%, old day 6:33.5 ± 1.8% , P>0.05) (Figure 3J). The spatial organization of macrophages and 3β-HSD-positive steroidogenic cells in organoids generated from old mice was similar to young counterparts in that macrophages were enriched in the perimeters of organoids and 3β-HSD-positive cells were excluded from the cores (Figure 3H, J).

### Age-dependent changes in actin and cell adhesion pathways in ovarian somatic cells are suggestive of increased cellular stiffness

To identify age-related changes in the ovarian somatic compartment at the cellular level, which may underlie altered organoid formation and function, we examined the gene expression patterns of primary ovarian somatic cells from reproductively young and old mice which were used as the input to generate the organoids (monolayer) (Figure 4A). This analysis revealed two additional clusters that were not present in the UMAP when cells dissociated from organoids were incorporated in the analysis (Figure 4A, Supplemental Figure S4A) ^5, 30, 31^. In the analysis of the monolayer alone, the stroma/mesenchyme cells segregated into two clusters, and myeloid cells and lymphocytes separated into distinct clusters (Figure 4A). To define the identity of cells present in each of the stroma/mesenchyme clusters we performed Gene Ontology analysis of genes differentially expressed within each cluster. High expression of *Col1a2* and enrichment of biological processes and cellular components related to ECM function in the stroma/mesenchyme 1 cluster suggested that cells in this cluster were matrix fibroblasts (Figure 4B, Supplemental Figure S4A) ^10^. Interestingly, pathways associated with the actin cytoskeleton were also enriched in this cluster (Figure 4B). To evaluate age-dependent changes in the stroma/mesenchyme 1 cluster, we performed Gene Ontology analysis of genes differentially expressed in this cluster with advanced reproductive age. Actin polymerization or depolymerization and actin filament organization pathways were upregulated with age, driven by increased expression of genes that play roles in actin binding, actin nucleation, actin capping, as well as adherens and tight junctions (Figure 4C-D, Table 1). Cell adhesion and cell migration pathways were downregulated with age (Figure 4C). Downregulation of cell adhesion-related pathways was driven by decreased expression of claudins, cadherins, and integrins, among other genes (Table 2). In general, increased F-actin content and organization are associated with increased cellular stiffness, which could contribute to decreased cell-cell adhesion ^33^. Of note, increased cellular stiffness is a hallmark of aging cells in other tissues, but to date has not been examined in the context of ovarian aging ^33^.

**Figure 4.**
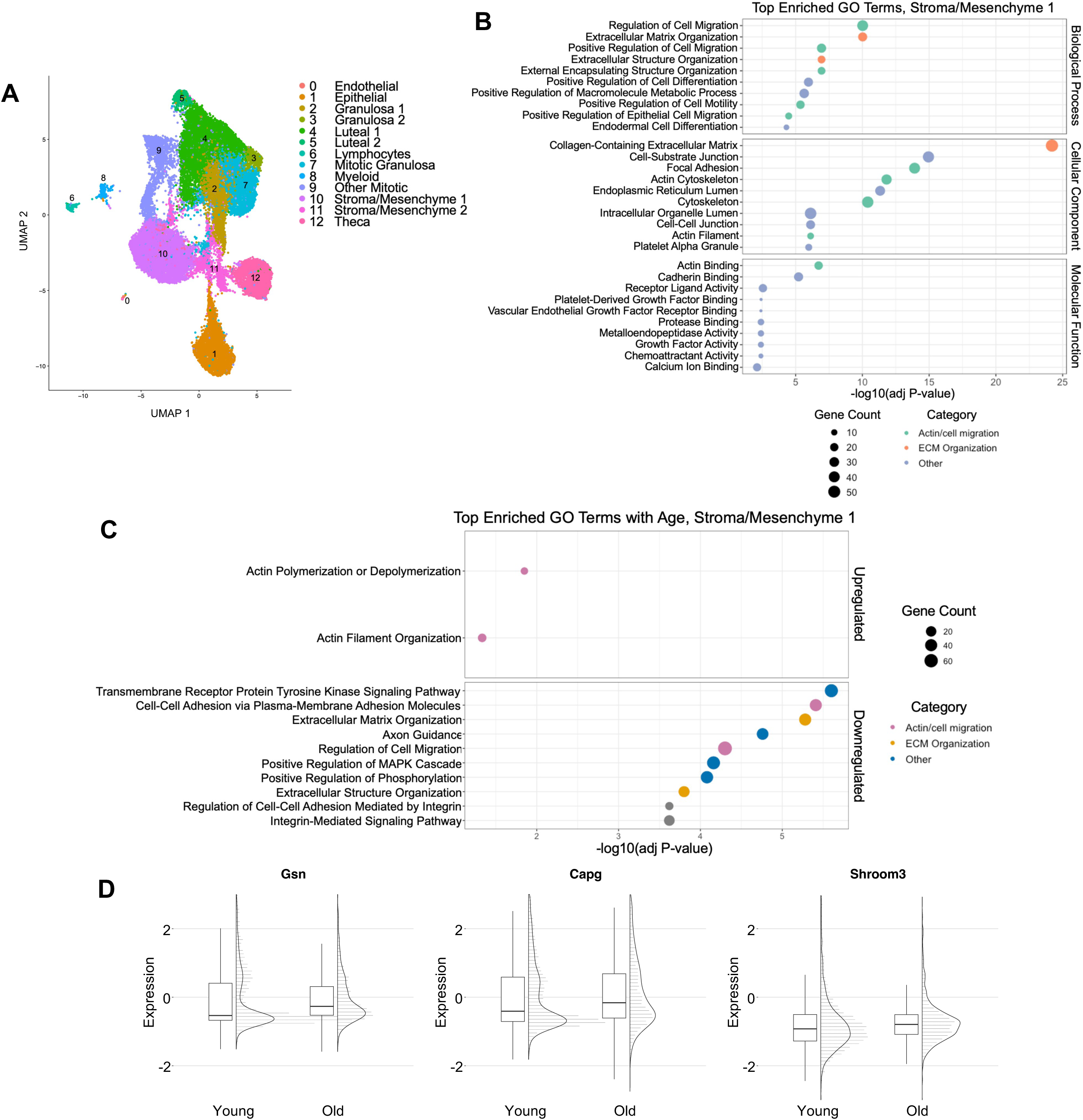
Actin pathways are upregulated in stroma/mesenchyme 1 cells with age and cell adhesion pathways are downregulated. A) Uniform manifold approximation and projection (UMAP) plot of only primary mouse ovarian somatic cells from reproductively young (6-12 weeks) and reproductively old (10-14 months) mice following plating in 2D, prior to organoid generation (monolayer). Unbiased clustering revealed 13 distinct cell populations. scRNAseq was performed for 2 replicates per group. B) Gene ontology (GO) analysis of differentially expressed genes within the Stroma/Mesenchyme 1 cluster compared to other cell clusters. Biological processes, cellular components, and molecular functions are labelled and displayed in manually grouped categories. C) GO analysis of differentially expressed genes in the Stroma/Mesenchyme 1 cluster with age for monolayer cells. Pathways upregulated and downregulated with age are labelled and displayed in manually grouped categories. D) Raincloud plots showing expression of genes driving upregulated actin-related pathways in the Stoma/Mesenchyme 1 cells from young and old mice.

**Table 1.** Genes driving upregulated actin pathways in stroma/mesenchyme 1 cells from reproductively old mice.

| Gene Symbol | Gene Name | Average Log2Fold-Change | Adjusted P-Value | Function of Encoded Protein<sup>1</sup> |
| --- | --- | --- | --- | --- |
| Lcp1 | Lymphocyte cytosolic protein 1 | 13.158772 | 3.05E-73 | Enables actin filament binding activity; Acts in actin filament bundle assembly |
| Gsn | Gelsolin | 10.3239145 | 1.57E-25 | Actin binding protein; Involved in actin filament capping; Involved in actin polymerization and depolymerization |
| Was | Wiskott-Aldrich syndrome | 5.30194898 | 2.99E-26 | Actin signaling; Actin nucleation and assembly |
| Shroom3 | Shroom family member 3 | 3.44015737 | 7.47E-10 | Enables actin binding activity |
| Capg | Capping actin protein, gelsolin like | 3.06743619 | 0.00053986 | Caps barbed ends of actin filaments |
| Avil | Advillin | 1.26971477 | 0.00056432 | Enables actin filament binding activity |
| Pstpip2 | Proline-serine-threonine phosphatase-interacting protein 2 | 1.1908554 | 5.52E-17 | Enables actin filament binding activity; Involved in actin filament polymerization |
| Aif1 | Allograft inflammatory factor 1 | 0.87640441 | 1.40E-22 | Enables actin filament binding activity |
| Cgnl1 | Cingulin-like 1 | 0.67256897 | 1.84E-10 | Mediates adherens and tight junction assembly and maintenance |

<sup>1</sup>Functions of encoded proteins were sourced from the NCBI Gene Database <sup>62</sup>.

**Table 2.**
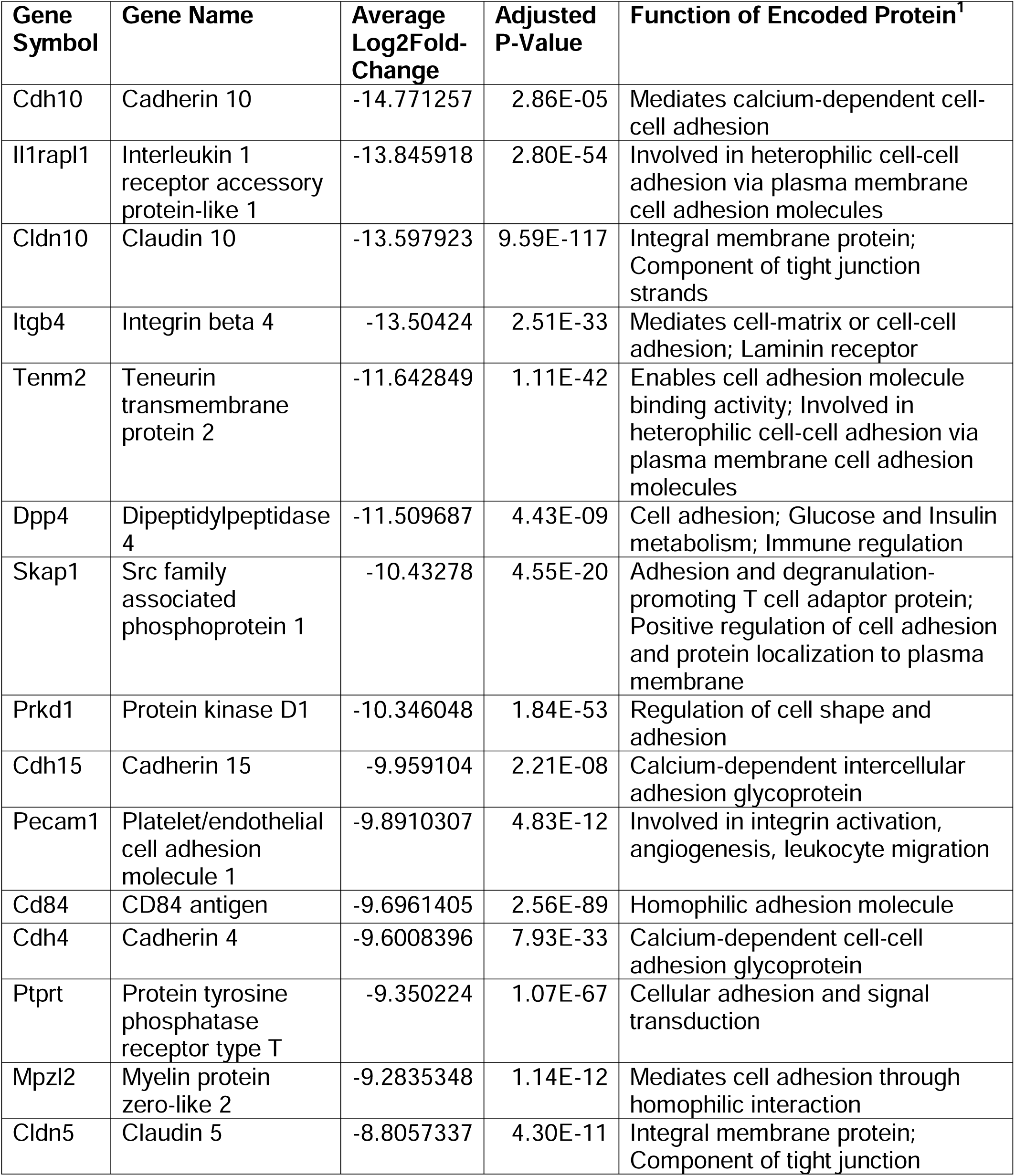
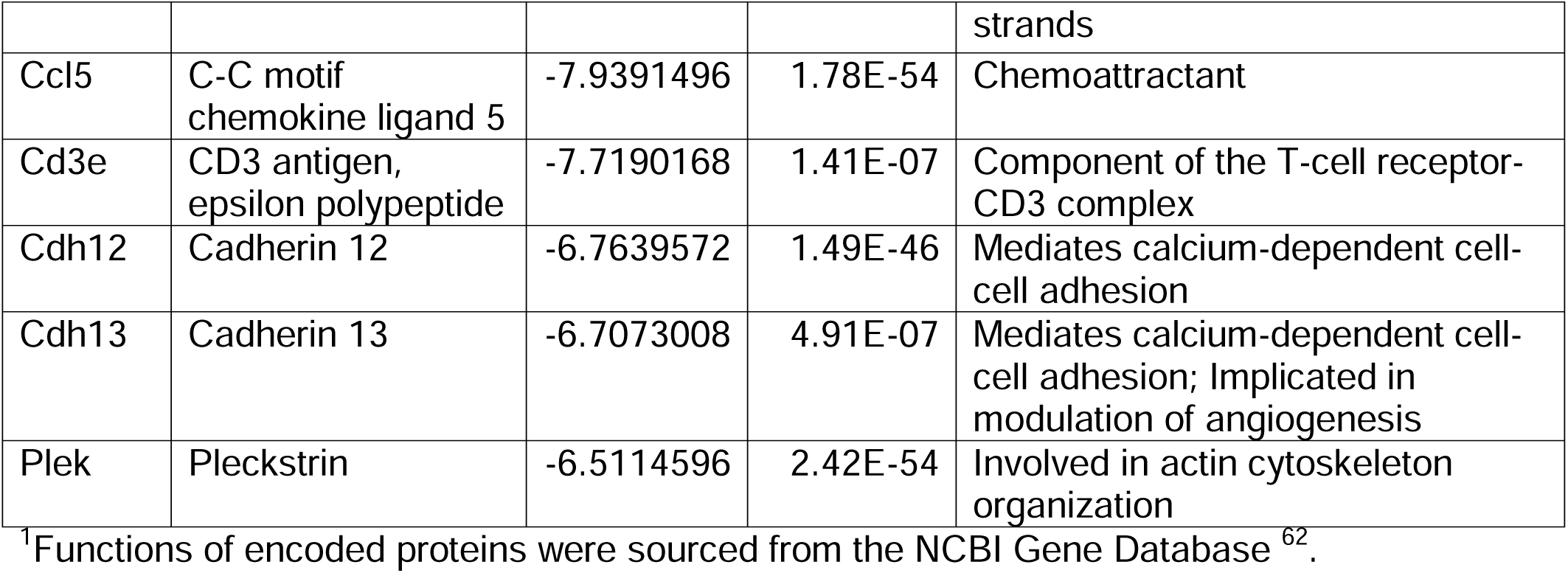
Top 20 genes greatest absolute Log_2_fold-change driving downregulated cell adhesion pathways in stroma/mesenchyme 1 cells from reproductively old mice.

### Ovarian somatic cells have heterogenous morphology that is altered with advanced reproductive age

In addition to impacting cellular stiffness, the actin cytoskeleton plays a central role in regulating cell morphology ^34, 35^. Cell morphology has been linked to biological properties of cells including cell cycle progression, metastatic potential, as well as gene expression ^36–38^. Importantly, cell morphology encodes aging information and can be used as a predictive biomarker of cellular age ^39^. To evaluate the morphologies of ovarian somatic cells isolated from reproductively young and old mice, we employed a previously established high throughput cellular phenotyping scheme which extracts hundreds of parameters describing the morphologies of cells and their corresponding nuclei ^36, 40^. These morphological parameters include metrics that describe both the sizes and shapes of cells and nuclei, thereby allowing direct interrogation and comparison of cell morphology at the single-cell level ^36, 40^. To this end, primary ovarian somatic cells isolated from young and old mice were fixed and stained to visualize F-actin, vimentin, and DNA (Figure 5A-B). Individual cells were segmented using CellProfiler to determine nuclear and cellular boundaries. From these boundaries we computed 228 morphological parameters along with the corresponding F-actin and vimentin expressions per cell (Figure 5A) ^36, 40^. These 228 parameters were subsequently reduced to 106 following orthogonality analysis to identify the parameters that best captured the variance within the dataset. Using these 106 parameters, we performed clustering analysis of more than 65,000 single cells collected from four biological replicates. (Figure 5A).

**Figure 5.**
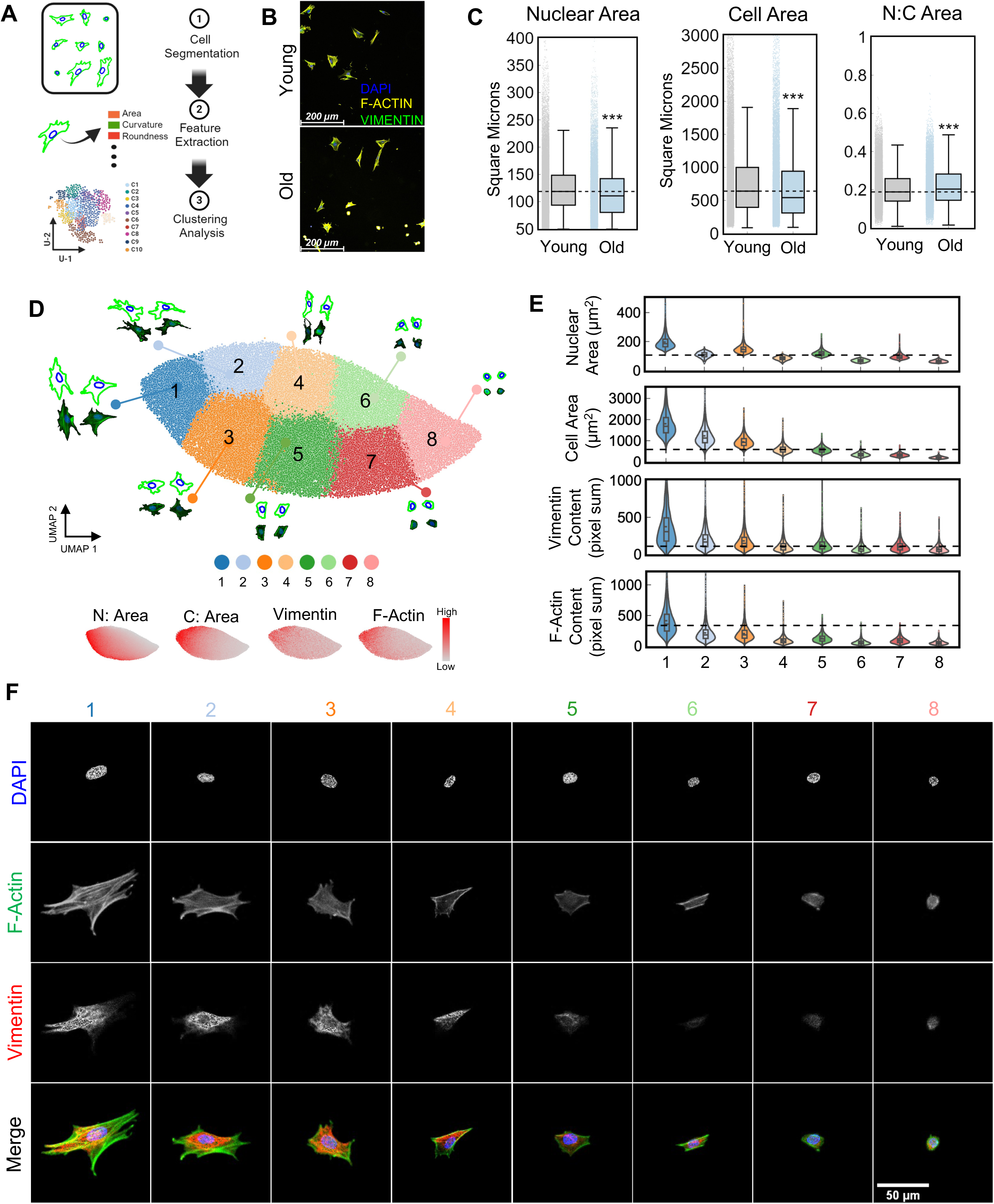
Ovarian somatic cells have heterogenous morphology. A) Schematic of the morphological analysis pipeline. B) Representative images of ovarian somatic cells from reproductively young (6-12 weeks) and reproductively old (10-14 months) mice. F-actin (phalloidin, yellow), vimentin (green), and DAPI (blue) were detected by immunocytochemistry. Scale bars = 200 µm. C) Quantification of nuclear area, cellular area, and nuclear to cellular area ratio for ovarian somatic cells from young and old mice. D) 2D UMAP visualization based on 106 morphological parameters for over 65,000 ovarian somatic cells. 8 k-means clusters are layered on top of the UMAP space with representative cellular (outer, green) and nuclear (inner, blue), and raw image morphologies for each cluster. Panels below show distribution of high (red) and low (gray) nuclear area (N: Area), cellular area (C: Area), vimentin expression, and F-actin expression displayed within the UMAP space. High and low represent standard scaling of expression across the dataset. E) Violin plots of the nuclear area, cellular area, vimentin, and F-actin content for each of the 8 k-means clusters. F) Representative images of an ovarian somatic cells within each morphological cluster. DAPI (blue), F-actin (phalloidin, green), and vimentin (red) were detected by immunocytochemistry. Brightness of fluorescent staining was adjusted equivalently for all images to highlight localization. Scale bar = 50 µm.

On average, ovarian somatic cells isolated from reproductively old mice had smaller nuclear and cellular area compared to young counterparts, resulting in a significant age-associated increase in the nuclear-to-cellular area ratio (Figure 5C). To gain deeper insights into the morphological differences, we performed clustering and dimensional reduction analyses of the cell and nuclear morphologies across conditions to identify groups of cells with similar morphologies. K-means clustering analysis visualized using 2D UMAP identified 8 morphological clusters of primary ovarian somatic cells, each with distinct cellular and nuclear morphologies (Figure 5D, F).

Cluster 1 corresponded to cells with the largest nuclear and cellular area, and highest F-actin and vimentin staining, suggesting a fibroblast identity (Figure 5D-F). Cluster 8, on the other hand, corresponded to the smallest and roundest cells with depleted F-actin and vimentin staining, suggestive of an epithelial identity (Figure 5D-F). In general, primary ovarian somatic cells isolated from old mice were enriched in cluster 8 compared to young counterparts (Figure 6A, D). The age-associated enrichment in this cluster is consistent with the increased abundance of ovarian epithelial cells with age (Figure 3C, Supplemental Figure S3A, C) ^17, 41^. Moreover, while cells from both young and old mice exhibited similar abundance of clusters 1, 2, 4, 6, and 7, cells from young mice were enriched in clusters 3 and 5 (Figure 6A, D). Interestingly, although the enriched morphological clusters were relatively consistent between different groups of young mice, ovarian somatic cell morphologies were highly heterogeneous among different groups of old mice (Supplemental Figure S5A). Taken together, we show that ovarian somatic cells have heterogenous morphology. Moreover, the relative abundance of various cell morphologies shifts with age, further demonstrating altered properties of the aging ovarian somatic compartment at a cellular level.

**Figure 6.**
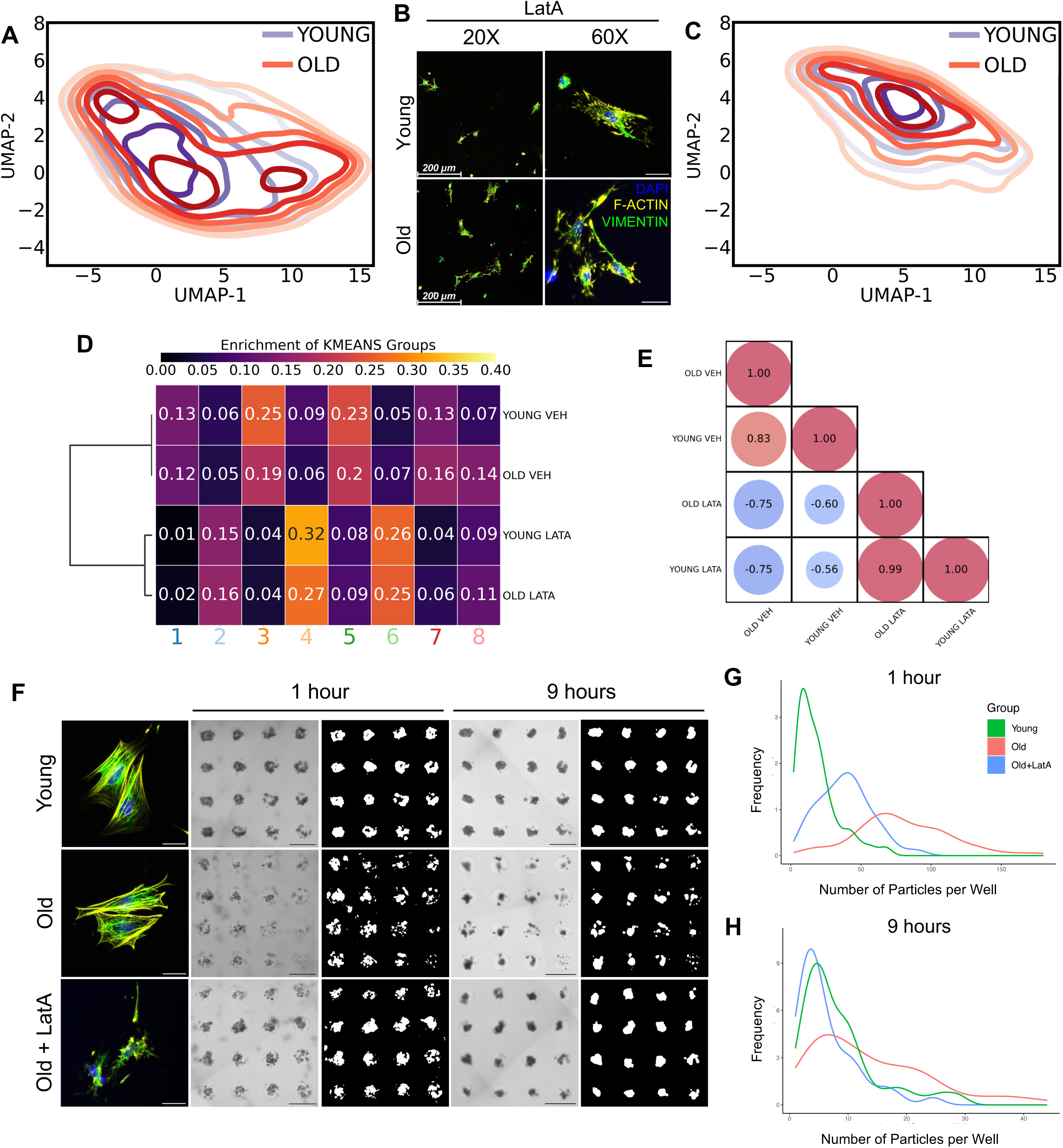
Disrupting the actin cytoskeleton shifts ovarian somatic cell morphology and improves the aggregation of ovarian somatic organoids generated from reproductively old mice. A) Contour overlays on the UMAP space showing enriched morphologies for ovarian somatic cells from young (purple) and old (red) mice. B) Representative 20X and 60X images of ovarian somatic cells from young and old mice treated with Latrunculin A (LatA). F-actin (phalloidin, yellow), vimentin (green), and nuclei (DAPI, blue) were detected by immunocytochemistry. 20X Scale bars = 200 µm. 60X Scale bars = 50 µm. C) Contour overlays on the UMAP space showing enriched morphologies for ovarian somatic cells from young (purple) and old (red) mice treated with LatA. D) Heatmap showing the fractional abundance of ovarian somatic cells from young and old mice with or without LatA treatment within each k-means cluster. E) Bubble plot showing correlation of morphological enrichment between experimental groups. F) Representative images of primary ovarian somatic cells following treatment with Latrunculin A (LatA) or vehicle control. F-actin (phalloidin, yellow), vimentin (green), and nuclei (DAPI, blue) were detected by immunocytochemistry. Scale bars = 50 µm. Representative images of organoids generated from young and old primary ovarian somatic cells following treatment with Latrunculin A (LatA) or vehicle control at 1 hour and 9 hours post seeding into agarose micromolds. Scale bars = 500 µm. Right panels for each timepoint are thresholded masks of cells in agarose micromolds. B-C) Kernel-smoothed distribution curves for histograms quantifying the frequency of different numbers of particles per well at 1 hour (G) and 9 hours (H) following seeding of primary ovarian somatic cells from young (green) and old (red) mice treated with a vehicle control or primary ovarian somatic cells from old mice treated with LatA (blue) into agarose micromolds. N=1 micromolds per condition. Representative images for N=2 additional micromolds per condition are shown in Supplemental Figure S6.

### Disrupting the actin cytoskeleton shifts ovarian somatic cell morphology and improves the aggregation of ovarian somatic organoids generated from reproductively old mice

Our results demonstrate an age-related upregulation of actin pathways in the stroma/mesenchyme 1 cluster and an age-dependent decrease in cell adhesion pathways, which is indicative of increased cellular stiffness (Figure 4C).Given the high actin content in cluster 1, we hypothesized that cells in this cluster may contribute to the age-dependent increase in cellular stiffness and that morphology of these cells may be responsive to actin depolymerization. To directly evaluate this, we treated primary cells isolated from reproductively young and old mice with 1 µM Latrunculin A (LatA) for 45 minutes, which depolymerizes actin filaments and sequesters actin monomers to prevent polymerization ^42^. Treatment with LatA successfully disrupted the actin cytoskeleton in primary ovarian somatic cells isolated from reproductively young and old mice (Figure 6B). Moreover, LatA treatment significantly depleted the abundance of cells in clusters 1 and 3, in addition to normalizing the age-associated enrichment in cluster 8 (Figure 6C-D). Additionally, LatA treatment shifted cell morphologies from both young and old mice from clusters 1, 3 and 5 to clusters 4 and 6, which are comparatively smaller cells with smaller nuclear area and reduced F-actin and vimentin content (Figure 5D-F, Figure 6C-D). Furthermore, ovarian somatic cells treated with Lat A exhibited homogeneous morphologies irrespective of the age of the mice from which they were isolated (Figure 6C-D). This homogeneity was evidenced by a correlation value of 0.99 when comparing the morphological enrichment of cells derived from reproductively young and old mice following LatA treatment, compared to a correlation value of 0.83 when comparing vehicle treated controls (Figure 6E).

Increased cellular stiffness may underlie the poor ability of ovarian somatic cells derived from old mice to aggregate into fully functional organoids (Figure 2A) ^33^. To evaluate the functional role of actin polymerization and cellular stiffness on age-dependent organoid formation, we tested whether partial depolymerization of the actin cytoskeleton, through a short-term treatment with LatA, could rescue the age-dependent impairment in organoid formation. Organoids generated using ovarian somatic cells from reproductively young mice served as a baseline control. Interestingly, LatA treatment of ovarian somatic cells derived from reproductively old mice resulted in enhanced organoid formation at 1 hour and 9 hours post cell seeding relative to vehicle treated controls (Figure 6F). In fact, quantitative analysis of the frequency of the number of particles per microwell demonstrated that LatA treatment of ovarian somatic cells isolated from reproductively old mice resulted in an organoid aggregation profile that more closely resembled that of young mice (Figure 6G-H; Supplemental Figure S6A-B). Overall, age-associated changes to the ovarian somatic compartment at a cellular level underlie differences in the capacity for aggregation.

## Discussion

In this study, we developed an innovative and robust organoid model of the mouse ovarian somatic compartment. This represents a major advance in the field in that it preserves the cellular heterogeneity of the native tissue, demonstrates organ-specific functionality, and provides novel insights into mechanisms of ovarian aging. Ovarian somatic organoids were solid structures with proliferative cells present throughout and apoptotic cells restricted to the perimeter. They contained key ovarian cell types, including fibroblasts, epithelial, endothelial, immune, granulosa, theca, and luteal cells, and self-organized such that 3β-HSD-positive steroidogenic cells were excluded from the core and macrophages localized to the perimeter. Additionally, organoids produced an extracellular matrix, were responsive to exogenous stimuli, and secreted hormones. Interestingly, organoids generated from reproductively old mice demonstrated impaired aggregation and decreased growth compared to young counterparts. Organoids exhibited age-dependent changes in relative cell composition and functional output with respect to endocrine function. At a cellular level, aging was associated with upregulation of pathways involved in actin dynamics in ovarian stroma/mesenchyme cells, as well as downregulation of pathways involved in cell adhesion. Enrichment of these pathways suggested increased cellular stiffness with age. Cell morphology, which is regulated by the actin cytoskeleton, was heterogenous among ovarian somatic cells and was altered with age. Modulation of actin with Latrunculin A shifted ovarian somatic cell morphology and improved aggregation of organoids from old mice. Overall, insights gained from this model suggest that aging of the ovarian somatic compartment may have cellular origins.

One of the greatest limitations of current *ex vivo* models of the ovarian somatic compartment is their inability to recapitulate and maintain cellular heterogeneity. Attempts at culturing ovarian somatic cells in 2D results in loss of heterogeneity over time in culture, and existing organoid models, including ovarian surface epithelial organoids, only capture a single cell type ^2, 21^. In contrast, we have demonstrated the maintenance of fibroblasts, epithelial, immune, and steroidogenic cell populations within organoids over six days in culture. Moreover, we have demonstrated functionality of these cell populations in their ability to respond to a pro-fibrotic stimulus as well as their ability to produce hormones. This fidelity in composition and function will allow use of this organoid model to study interactions between the ovarian stroma and other cells and tissues. Importantly, we demonstrate that oocytes and intact follicles are not needed to assemble and organize the ovarian stroma, but future studies may use ovarian somatic organoids to understand the impact of oocyte-secreted factors on the stroma and vice versa, particularly in the context of aging and the corresponding decline in oocyte quality and quantity ^43^. Furthermore, these organoids can be generated using rhesus macaque ovarian tissue, enabling translation of this model for primate studies. In addition to applying this model to other species, these methods may also be used to generate organoids to probe the contribution of the ovarian stroma in conditions such as polycystic ovary syndrome, ovarian cancer, and post-chemotherapy or radiation.

Ovarian somatic organoids exhibited several age-dependent changes, including differences in relative cell composition. Organoids generated from reproductively old mice had a larger proportion of epithelial cells than young counterparts, which may be due to ovarian surface epithelial hyperplasia that occurs with age ^17, 41, 44, 45^. Consistent with the age-associated increase in epithelial cells, ovarian somatic cells from reproductively old mice were enriched for a population of cells with small, rounded morphology and low vimentin expression, suggesting epithelial identity. Moreover, organoids from old mice contained fewer steroidogenic cells, especially granulosa cells, which is consistent with age-dependent depletion of ovarian follicles ^46^.

Correspondingly, organoids from old mice exhibited blunted hormone production compared to organoids from young mice. This attenuated hormone secretion could be due to the smaller number of steroidogenic cells in addition to reduced functionality of the remaining cells, given the downregulation of pathways related to oxidative stress response in luteal 2 cells with age. Additionally, although present at low abundance, organoids from old mice had an increased proportion of immune cells, reflective of the inflammatory environment of the aging ovary ^5, 6, 47^.

Organoids did not recapitulate age-related changes to the extracellular matrix observed *in vivo*, particularly the age-dependent increase in collagen and decrease in hyaluronan ^6, 11^. Myofibroblasts are the primary cell type responsible for ECM secretion during wound healing and fibrosis ^6, 48^. Thus, the inability of organoids to establish the age-related increase in collagen content present in the ovary *in vivo* may be due, in part, to the similar percentage of myofibroblasts in organoids derived from both young and old mice at day 1 of culture. It is possible that the procedure of isolating of primary ovarian somatic cells from reproductively old mice selects for the healthiest cells, particularly due to age-associated ovarian fibrosis. In fact, isolation of hepatic stellate cells, the main fibrogenic cell type in the liver, from fibrotic tissue is more difficult than isolation from normal tissue and requires a modified protocol ^49^. Additional studies may be required to optimize isolation of activated myofibroblasts from reproductively old ovaries, as well as to determine whether aging resets when cells are removed from an old ovary or whether aging phenotypes persist *in vitro* over an extended period of time. Interestingly, organoids from old mice contained a significantly higher proportion of myofibroblasts by day 6 of culture, suggesting that age-related changes may in fact re-establish *in vitro*. Given the time required for matrix deposition and remodeling, a longer time in culture may be needed to observe age-associated changes in the ECM. Nevertheless, several robust age-dependent differences are captured in the organoid model, including the striking contrast in aggregation potential of organoids from reproductively young and old mice, which may be a result of an increase in cellular stiffness with advanced reproductive age.

In the context of ovarian aging, to date “stiffness” has referred to altered biomechanical properties at the level of the tissue due to age-associated changes in the ECM ^6, 10–13^. However, aging is also associated with increased stiffness at a cellular level ^33^. Studies using atomic force microscopy to evaluate stiffness of human epithelial cells and fibroblasts have demonstrated that increased force is required to indent these cells with age ^50, 51^. An age-associated increase in cell stiffness is correlated to changes in the cytoskeleton, in particular increased levels of F-actin, as well as increased cytoskeletal crosslinking and bundling ^52–54^. Age-dependent changes to cell mechanics may contribute to functional decline given mechanical regulation of cell migration and adhesion ^33^. Cell stiffness has been examined in relation to ovarian cancer and those studies demonstrated that reduced cell stiffness can be utilized as a biomarker of metastatic potential, as malignant cells are softer than non-malignant counterparts ^55^. However, additional studies are required to further probe cellular stiffness and mechanics in distinct morphological populations of ovarian somatic cells, as well as to determine how they contribute to the progression of reproductive aging.

In our study, we demonstrated upregulation of actin polymerization or depolymerization and actin filament organization pathways in ovarian stroma/mesenchyme cells with advanced age, in addition to the presence of morphologically large cells with high actin content. This age-dependent increase in actin content and organization pathways is coupled with downregulation of cell adhesion and cell migration pathways, as well as phenotypically reduced aggregation of ovarian somatic cells into organoids. These findings are suggestive of an age-dependent increase in cell stiffness of ovarian somatic cells. Consistent with this, depolymerization of actin using Latrunculin A shifted ovarian somatic cell morphology and resulted in a partial rescue of organoid formation. Further studies are required to determine if a specific population of ovarian somatic cells is exhibiting an age-associated increase in cellular stiffness. Future research may also seek to determine if cellular stiffness is a consequence of or a contributing factor to the aging ovarian microenvironment.

Overall, we have developed a completely novel and robust model system of the mammalian ovarian somatic compartment, which maintains the complex cellular heterogeneity, cellular interactions, and functions seen *in vivo*. This organoid model can be utilized to interrogate ovarian aging and identify new mechanisms underlying this process. Using this model in addition to single cell transcriptomic and morphological analyses, we have uncovered cellular stiffness as a mechanism potentially underlying ovarian aging for future studies.

## Methods

### Animals

Female CD-1 mice aged 5 weeks and female CD-1 retired breeders were purchased from Envigo (Indianapolis, IN). Upon arrival to Northwestern University, mice were acclimated for at least 1 week and subsequently used for experiments at 6-12 weeks (reproductively young) or 10-14 months (reproductively old). Mice were housed in a controlled barrier facility at Northwestern University’s Center for Comparative Medicine (Chicago, IL) under constant temperature, humidity, and light (14h light/10h dark). Upon arrival to Northwestern University, animals were provided water and Teklad Global irradiated chow (2916) containing minimal phytoestrogens *ad libitum*. All mouse experiments were performed under protocols approved by the Institutional Animal Care and Use Committee (Northwestern University) and in accordance with the National Institutes of Health Guide for the Care and Use of Laboratory Animals.

Rhesus macaque (Age: 18 years) ovarian tissue obtained following necropsy was cut into quarters and submerged in SAGE OTC Holding Media (Cooper Surgical, Trumbull, CT) in 5 mL tubes. Tubes were shipped from Beaverton, OR to Chicago, IL with ice packs to maintain the temperature at 4 °C. Upon arrival in Chicago, a piece of tissue was fixed in Modified Davidson’s (Electron Microscopy Sciences, Hatfield PA) overnight at 4°C and the remaining tissue was processed for somatic cell isolation. The time between surgery and tissue processing for somatic cell isolation was approximately 20 hours. The general care, housing, and use of rhesus macaques for experiments was performed under research protocols approved by the Oregon National Primate Research Center (ONPRC) institutional animal care and use committee.

### Ovarian somatic cell isolation

Primary murine ovarian somatic cell isolation and plating were performed as previously described ^2, 56^. Briefly, mouse ovaries were harvested, separated from the bursa, and placed in Dissection Media composed of Lebovitz’s Medium (L15, Gibco, Grand Island, NY) supplemented with 1% fetal bovine serum (FBS, Gibco, Grand Island, NY) and 0.5% Penicillin-Streptomycin (Gibco, Grand Island, NY). To enrich for the stromal fraction, antral follicles were mechanically punctured using insulin syringes to release cumulus-oocyte-complexes. The remaining stroma-enriched fraction was dissected into smaller pieces using forceps and incubated in _a_MEM Glutamax (Gibco, Grand Island, NY) supplemented with 1% FBS, 0.5% Penicillin-Streptomycin, 82 units/mL Collagenase IV (ThermoFisher Scientific, Waltham, MA), and 0.2 mg/mL DNase I (ThermoFisher Scientific, Waltham, MA) for 30 minutes at 37°C in a humidified environment of 5% CO_2_ in air. To assist with enzymatic digestion, the tissue was triturated every 15 minutes by pipetting. After 30 minutes, the enzymatic digestion was quenched with an equal volume of _a_MEM Glutamax supplemented with 10% FBS. The cell suspension was then filtered through a 40 µm strainer, pelleted (300 x g for 5 minutes at room temperature), washed using RPMI 1640 containing 25 mM HEPES and 2 mM L-glutamine (Gibco, Grand Island, NY) supplemented with 10% FBS and 1% Penicillin-Streptomycin (Plating Media), and plated. Following plating overnight, cells were washed with Dulbecco’s PBS without calcium or magnesium (DPBS, Gibco, Grand Island, NY) and lifted from 2D culture using 0.05% Trypsin-EDTA (Gibco, Grand Island, NY).

For cell morphology analysis, slides were coated 50 µg/mL collagen I (Corning, Corning, NY) diluted in DPBS for 90 minutes at 37°C. Cells were strained through a 40 µm cell strainer and seeded onto collagen-coated slides at a density of 58,000 cells/slide. Following culture on collagen-coated slides overnight, cells were treated with 1 µM Latrunculin A (LatA, Sigma-Aldrich, St. Louis, MO) or vehicle (DMSO, Sigma-Aldrich, St. Louis, MO) and incubated at 37°C for 45 minutes, as noted in figure legends. Slides were then washed with DPBS and fixed with 4% Paraformaldehyde (Electron Microscopy Sciences, Hatfield, PA) for 12 minutes at room temperature.

For isolation of primary somatic cells from rhesus macaque ovarian tissue, ovarian tissue quarters were cut into 500 µm thick slices using a Stadie-Riggs Tissue Slicer to separate the ovarian cortex and medulla. Tissue slices were placed in Dissection Media and cut into pieces 2-3 mm x 2-3 mm using a scalpel. Tissue pieces were transferred to _a_MEM Glutamax supplemented with 0.1% (w/v) bovine serum albumin (BSA, Sigma Aldrich, St. Louis, MO), 1X Insulin-Transferrin-Selenium (ITS, Sigma Aldrich, St. Louis, MO), 40 µg/mL Liberase DH (Sigma Aldrich, St. Louis, MO), 82 units/mL Collagenase IV, and 0.2 mg/mL DNase I and incubated for 1 hour at 37°C in a humidified environment of 5% CO_2_ in air. To assist with enzymatic digestion, the tissue was triturated every 15 minutes by pipetting. After 1 hour, the digestion was quenched with sterile filtered FBS. The cell suspension was then filtered through a 40 µm strainer, pelleted (300 x g for 5 minutes at room temperature), washed using DMEM/F12 (Gibco, Grand Island, NY) supplemented with 10% FBS and 1% Penicillin-Streptomycin, and plated. Following plating overnight, cells were washed with PBS and lifted from 2D culture using 0.05% Trypsin-EDTA.

### Organoid generation and culture

Organoids were generated using the MicroTissues 3D Petri Dish scaffold-free, 3D cell culture system (Sigma-Aldrich, St. Louis, MO, Z764043). Micromolds were cast using 1.5% Agarose (Hoefer Inc, Holliston, MA) dissolved in DPBS and stored at 4°C in DPBS with 1% Penicillin-Streptomycin until use. Agarose micromolds were pre-equilibrated in 500 µL IntestiCult (StemCell Technologies, Vancouver, CA) at 37°C prior to cell seeding. Following trypsinization, cells were resuspended in IntestiCult at a density of 250,000 cells per 75 µL media and 75 µL cell suspension was seeded into agarose micromolds. An additional 250 µL IntestiCult was added surrounding micromolds at least 1 hour after cell seeding. For data shown in Figure 1, HepatiCult (StemCell Technologies, Vancouver, CA), MesenCult (StemCell Technologies, Vancouver, CA), or Plating Media was used to pre-equilibrate agarose micromolds and resuspend cells as indicated. Media was changed the day following cell seeding and every other day throughout culture.

For TGF-β treatment, culture media surrounding agarose micromolds was replaced with IntestiCult containing 10 ng/mL TGF-β (R&D Systems, Minneapolis, MN) on day 3 of culture and organoids were cultured for an additional 48 hours. Following treatment, organoids were harvested from micromolds, pelleted by centrifugation (500 x g for 10 minutes at 4°C), and frozen on dry ice for RNA extraction.

For Latrunculin A treatment, following trypsinization, cells were resuspended in Plating Media containing 1 µM Latrunculin A or vehicle and incubated at 37°C for 30-45 minutes. Latrunculin A treatment was performed in polypropylene tubes to prevent cell adhesion. Following treatment, cells were pelleted, resuspended in IntestiCult, and seeded into agarose micromolds to assess organoid formation.

### Tissue processing, histology and immunohistochemistry

For histology and immunohistochemistry, micromolds were sealed with agarose. Sealed agarose micromolds and mouse ovaries were fixed in Modified Davidson’s overnight at 4°C. Following fixation, samples were transferred to 70% ethanol and stored at 4°C until processing. Samples were dehydrated using an automated tissue processor (Leica Biosystems, Buffalo Grove, IL), embedded in paraffin, and sectioned (5 µm thickness).

Hematoxylin & Eosin staining was performed following a standard protocol. Tissue sections were cleared with Citrosolv (Fisher Scientific Pittsburgh, PA) in 3, 5-minute incubations and mounted with Cytoseal XYL.

Picrosirius Red (PSR) staining was performed as previously published ^6, 11^. Briefly, tissue sections were deparaffinized in Citrosolv, rehydrated in graded ethanol baths (100, 70, and 30%) and washed in RO water. Slides were immersed in PSR staining solution for 40 minutes, then incubated in acidified water (0.05M hydrochloric acid) for 90 seconds. Tissue sections were dehydrated in 100% ethanol, cleared in Citrosolv, and mounted with Cytoseal XYL.

Hyaluronan-binding protein (HABP) assays were performed as previously published ^11, 57^. Briefly, tissue sections were deparaffinized in Citrosolv, rehydrated in graded ethanol baths (100, 95, 85, 70, and 50%), and washed in RO water.

Endogenous Avidin and Biotin were blocked using an avidin/biotin blocking kit (Vector Laboratories, Burlingame, CA) and non-specific antigens were blocked with 10% normal goat serum (Vector Laboratories, Burlingame, CA). Slides were then incubated with biotinylated-HABP (Calbiochem, San Diego, CA) for 1 hour at room temperature. Signal amplification was performed using the Vectastain Elite ABC Kit (Vector Laboratories) and subsequently the TSA (Tyramide Signal Amplification) Plus Fluorescein Kit (1:400; Akoya Biosciences, Marlborough, MA). Slides were mounted with Vectashield containing 4′,6-diamidino-2-phenylindole (DAPI, Vector Laboratories, Burlingame, CA). Slides treated with 1 mg/mL hyaluronidase (Sigma-Aldrich, St. Louis, MO) were utilized as negative controls.

For chromogenic immunohistochemistry (IHC), tissue sections were deparaffinized in Citrosolv, rehydrated in graded ethanol baths (100, 95, 85, 70, and 50%), and washed in RO water. Antigen retrieval was performed by microwaving slides in 1X Reveal Decloaker (Biocare Medical, Concord, CA) at 50% power for two minutes followed by 10% power for seven minutes. Endogenous peroxidase activity was blocked with 3% hydrogen peroxide, endogenous avidin and biotin were blocked using an avidin/biotin blocking kit, and non-specific antigens were blocked with 10% normal goat serum and 3% BSA in Tris-buffered saline (TBS). Slides were then incubated with the respective primary antibody (Supplemental Table S2) diluted in TBS containing 3% BSA in a humidified chamber overnight at 4°C. Slides were washed in TBS containing 0.1% Tween-20 (Sigma Aldrich, St. Louis, MO) (TBS-T) and incubated in secondary antibody (Supplemental Table S2) for 1 hour at room temperature. Signal amplification was performed using the Vectastain Elite ABC Kit and detection was performed using 3,3’-diaminobenzidine (DAB) with the DAB Peroxidase (HRP) Substrate Kit (Vector Laboratories, Burlingame, CA). Tissue sections were counterstained with hematoxylin, dehydrated in graded ethanol baths (80, 95, 100%), cleared in Citrosolv, and mounted with Cytoseal XYL. For immunofluorescent IHC, tissue sections were deparaffinized, rehydrated, and antigen retrieval was performed as described above. Non-specific antigens were blocked with 10% normal goat serum in TBS containing 0.3% Triton X-100 (Alfa Aesar, Haverhill, MA). Slides were then incubated with the respective primary antibody (Supplemental Table S2) in a humidified chamber overnight at 4°C. Slides were washed in TBS-T and incubated in secondary antibody (Supplemental Table S2) for 1 hour and 45 minutes at room temperature. Slides were subsequently washed in TBS-T and signal amplification was performed using the Vectastain Elite ABC Kit and TSA (Tyramide Signal Amplification) Plus Fluorescein Kit. Slides were mounted with Vectashield containing DAPI.

### Immunocytochemistry

Following fixation of cells seeded on collagen-coated slides, slides were washed 3 times with DPBS and cells were permeabilized with 0.01% Triton X-100 in DPBS for 12 minutes. Slides were blocked with 0.1% BSA in DPBS for 45 minutes and then incubated for 1 hour at room temperature with a primary antibody against vimentin (Supplemental Table S2) diluted in 0.1% BSA in DPBS. Slides were then incubated for 1 hour at room temperature, protected from light, with goat anti-rabbit Alexa Fluor 488 (Supplemental Table S2) and either rhodamine phalloidin (1:50, Invitrogen, Carlsbad, CA) or Alexa Fluor 633 phalloidin (1:50, Invitrogen, Carlsbad, CA) and CellMask Orange plasma membrane stain (1:10000, H32713, Invitrogen, Carlsbad, CA) diluted in 0.1% BSA and mounted in Vectashield containing DAPI.

### Dissociation for scRNA-seq

To process primary ovarian somatic cells (monolayer) for scRNAseq, cells were strained through a 40 µm cell strainer (PluriSelect, El Cajon, CA) following trypsinization, pelleted at 500 x g for 5 minutes, and resuspended in 100 µL DPBS containing 1% BSA. Organoids were harvested from agarose micromolds and dissociated in 0.25% trypsin (Gibco, Grand Island, NY) at 37°C in ultra-low attachment plates for 1 hour while pipetting every 10 minutes. Trypsin was quenched with Plating Media, cells were strained through a 40 µm cell strainer, pelleted at 500 x g for 5 minutes, washed with DPBS containing 1% BSA, and resuspended in 100 µL DPBS containing 1% BSA.

### scRNA-seq library construction and sequencing

Cell number and viability were analyzed using Nexcelom Cellometer Auto2000 using the AOPI fluorescent staining method. Sixteen thousand cells were loaded into the Chromium iX Controller (10X Genomics, Pleasanton, CA, PN-1000328) on a Chromium Next GEM Chip G (10X Genomics, Pleasanton, CA, PN-1000120), and processed to generate single cell gel beads in the emulsion (GEM) according to the manufacturer’s protocol. The cDNA and library were generated using the Chromium Next GEM Single Cell 3’ Reagent Kits v3.1 (10X Genomics, Pleasanton, CA, PN-1000286) and Dual Index Kit TT Set A (10X Genomics, Pleasanton, CA, PN-1000215) according to the manufacturer’s manual. Quality control for constructed library was performed by Agilent Bioanalyzer High Sensitivity DNA kit (Agilent Technologies, Santa Clara, CA, 5067-4626) and Qubit DNA HS assay kit for qualitative and quantitative analysis, respectively. The multiplexed libraries were pooled and sequenced on Illumina Novaseq6000 sequencer with 100 cycle kits using the following read length: 28 bp Read1 for cell barcode and UMI, and 90 bp Read2 for transcript.

### scRNA-seq quality control and data analysis

FASTQ files were generated and demultiplexed from base call (.bcl) files using Cell Ranger from 10X Genomics. Cell Ranger was also used for alignment of the FASTQ files to the mouse reference genome (mm10) and to count the number of reads from each cell that align to each gene. Resulting matrix files were imported in Seurat (v.4.0) and individual samples were pre-processed, normalized, and scaled ^58^. Cells with greater than 7% mitochondrial gene expression, between 500-6,000 features (unique genes), less than 600 or more than 50,000 counts (UMIs), less than 19% of all counts from ribosomal markers, and greater than 0.5% of all counts from red blood cell markers were removed. All samples were combined into a single dataset, with additional metadata containing original sample information. Clusters were separated at the 0.3 resolution using the FindClusters function in Seurat and differential gene expression analysis was performed to identify biomarkers to define cell clusters and compare between samples. Cell identities were assigned manually based on biomarker genes expressed in each cluster and endothelial and immune clusters were manually separated. A second dataset was created only using cells from reproductively young and old mice following plating in 2D, but prior to organoid generation (monolayer).

Clusters were separated at the 0.4 resolution and cell identities were assigned manually based on biomarker genes expressed in each cluster. All UMAPs, violin, and raincloud plots were created using tools in Seurat. The CellChat package in R was utilized for cell-cell communication analysis. All gene ontology analysis was performed using Enrichr with differentially expressed genes with adjusted P-values < 0.05 and absolute log_2_ fold-change > 0.58 ^59^. Dot plots were created using the ggplot2 package in R to visualize enriched GO pathways.

### RNA extraction and real-time quantitative PCR (RT-qPCR)

Total RNA was isolated using a RNeasy Mini Kit (Qiagen, Germantown, MD) with on-column DNA removal using a RNase-Free DNase Set (Qiagen, Germantown, MD) following the manufacturer’s protocols. A SuperScript™ III First-Strand Synthesis SuperMix kit (Thermo Scientific, Waltham, MA) was used to generate cDNA following the manufacturer’s protocol. Real-time quantitative PCR (RT-qPCR) was performed using PowerUp SYBR Green Master Mix (Applied Biosystems, Thermo Scientific, Waltham, MA) on an Applied Biosystems QuantStudio 7 Pro real-time PCR machine (Applied Biosystems, Thermo Scientific, Waltham, MA). Melt curves were performed to confirm the presence of a single product per primer pair and positive and negative controls confirmed the quality of the run. Primer sequences were accessed from PrimerBank and are listed in Supplemental Table S3 ^60^. Relative fold-change of target genes between control and treated samples was determined using the 2^−ΔΔCt^ method ^61^.

### Enzyme-linked immunosorbent assays (ELISAs)

750 µL conditioned media from organoids generated from reproductively young and old mice was collected at days 1, 3, and 5 of culture and replaced with 750 µL fresh media. The concentrations of estradiol (E2) and progesterone (P4) in the conditioned culture media were measured using the Estradiol ELISA Kit (Cayman Chemical, Ann Arbor, MI) and the Progesterone ELISA Kit (Cayman Chemical, Ann Arbor, MI), respectively. Assays were performed according to the manufacturer’s instructions.

### Image segmentation and quantification for cell morphology analysis

Fluorescent images of cells from all conditions were acquired with a Leica Stellaris 5 Confocal Microscope at 20X magnification using 3 laser lines (405 Diode, 488 Diode, 568 Diode). Images were taken at 1024 x 1024-pixel resolution at 0.568 microns per pixel. CellProfiler^TM^ software was used to segment cells and nuclei from raw fluorescent TIFF images (DAPI was used to identify nuclei and phalloidin or CellMask plasma membrane stain was used to identify cellular structure, particularly in the Lat A treated cells with depleted actin structure). Subsequently, in-house curation pipelines were used to remove areas of high confluency and ensure single, well segmented cells for analysis. The curated masks were leveraged to quantify vimentin and actin content per cell. Specifically, the summation of pixel intensities of fluorescently stained vimentin and actin were quantified per cell. To assess F-actin texture, the *InfoMeas* parameter of the CellProfiler^TM^ texture measurement module was analyzed.

Over 65,000 single cells spanning both ages and treatments (vehicle and LatA) served as the experimental basis for the morphological analysis.

### Morphological feature extraction

228 morphological parameters were quantified per cell from individual nuclear and cellular masks (features extracted from in-house pipeline). These morphological parameters ranged from basic geometric features such as area and perimeter to more complex features such as roughness and curvature. To identify key parameters driving variance in the cell sample set, 2D primary factor analysis was performed across all cells and all morphological parameters. 2D primary factor was selected as it would fall in line closer to the subsequent, nonlinear 2D UMAP reduction conducted in the paper for selected parameters. Parameters expressing a resulting communality value below 0.5 were dropped for subsequent analysis to clean noise in the dataset (note a higher communality value indicates a parameter capturing more of the variance in a population), resulting in 106 morphological features retained for analysis.

### UMAP and k-means morphological analysis

To compress the 106-vector morphological space to a more interpretable format, a 2D UMAP transformation was created (reducing the larger vector space to two orthogonal vectors). UMAP is a nonlinear dimensionality reduction algorithm that seeks to capture the structure of high dimensional data in a lower-dimensionality space that keeps spatial relations intact. The morphological dataset was first cleaned through a log-transformation and standard scaling (scaling each variable between -1 to 1 and removing magnitude effects across variables). A UMAP model was trained on the preprocessed dataset, creating two constituent vectors (UMAP-1 and UMAP-2) with values for each individual cell, allowing for easily interpretable data visualization. Each point in the UMAP space represents an individual cell whose morphological parameters have been transformed and projected. In addition to UMAP, k-means clustering, an unsupervised hierarchical clustering method, was used to discretize distinct morphological groups within the preprocessed dataset. An optimal number of clusters, 8, was calculated by a plateau in the inertia and silhouette values of the k-means algorithm. The algorithm was applied to each individual cell, resulting in single cells with unique UMAP-1, UMAP-2, and k-means coordinates.

### Imaging and image analysis

Transmitted light images of organoids were taken with an EVOS FL Auto Cell Imaging System (ThermoFisher Scientific, Waltham, MA) using a 4X or 10X objective. To view entire agarose micromolds, scans comprised of a series of individual images were taken across the micromold and then automatically stitched together using the EVOS software. Images of tissue sections following histological and chromogenic immunohistochemical staining were taken with a Nikon Eclipse E600 epifluorescence microscope (Nikon, Melville, NY) using a 63X objective, a SPOT Insight 2.0 Megapixel color camera (SPOT Imaging, Diagnostic Instruments, Sterling Heights, MI), and the associated SPOT software (SPOT Imaging, Diagnostic Instruments, Sterling Heights, MI). Images of organoid tissue sections following HABP assays and immunofluorescent immunohistochemical staining were taken with a Leica SP5 inverted laser scanning confocal microscope (Leica Microsystems, Buffalo Grove, IL) using near-UV (405 nm) and argon (488 nm) lasers. Settings were determined by the Glow-Over Look Up Table (LUT) using the LAS AF software (Leica Microsystems, Buffalo Grove, IL) to ensure that there was no pixel oversaturation. Epifluorescence images of ovarian tissue sections were taken with an EVOS FL Auto Cell Imaging System equipped with DAPI (Excitation 357/44 nm, Emission 447/60 nm) and GFP (Excitation 470/22 nm, Emission 510/42 nm) LED light cubes using a 20X objective. Imaging settings, including light, gain, and exposure times, were kept consistent between samples.

FIJI software (National Institutes of Health, Bethesda, MD) was used to quantify average area per organoid and average number of organoids per microwell from transmitted light scans. Thresholds were adjusted for each image to capture all cells and the Analyze Particles function was used to count the number and area of thresholded particles. To quantify the average area per microwell, the total thresholded area was divided by the number of visible microwells. Particles larger than 300 µm^2^ were considered organoids and used for analysis. To quantify the average number of organoids per microwell, the total number of organoids was divided by the number of visible microwells. Organoid area following removal from agarose micromolds was measured manually using the freehand tool in FIJI.

Image analysis following histological and immunohistochemical staining was performed using FIJI software. Total organoid area was measured manually using the freehand tool. To measure positive staining, color deconvolution was performed to isolate the DAB staining from images following chromogenic IHC or the channel of interest was separated from images following immunofluorescent IHC and HABP assays. Thresholds were set based on the highest intensity signal for each marker and kept consistent for all organoids. To calculate percent positive area, thresholded area was divided by the total area for each organoid. Four representative regions of interest (ROIs) in the ovarian stroma and corpora lutea were analyzed for each marker. Total area of ROIs was determined by thresholding and thresholds to capture positive staining were set as described above.

To quantify the number of particles per well following vehicle or LatA treatment, ROIs were drawn over each microwell in transmitted light scans. The total number of particles was measured using the Analyze Particles function following thresholding of each ROI. ROIs containing a majority of cells not within the plane of focus were excluded from analysis.

## Statistical Analysis

Statistical analysis was performed using GraphPad Prism Software (La Jolla, CA) or R (RStudio, version 4.3.0). T-tests, one-way ANOVAs, two-way ANOVAs (mixed-effects analysis) followed by Tukey’s post hoc test for multiple comparisons, or nonparametric Kruskal-Wallis tests followed by Dunn’s multiple comparisons tests were performed to evaluate differences between experimental groups as indicated in the figure legends. P-values <0.05 were considered significant. All experiments were performed with at least two independent cohorts of at least five mice and all total organoid or replicate numbers are specified in the figure legends.

## Acknowledgements

This work was supported by the National Institute of Child Health and Human Development (R01HD093726 to F.E.D. and M.T.P, T32HD094699 to S.S.D. and E.J.Z.), the National Cancer Institute (F31CA257300 to S.S.D.), startup funds from the Department of Obstetrics and Gynecology (to F.E.D.), and P51 OD011092 (DPCPSI, ORIP, NIH) to ONPRC. We would like to acknowledge Dr. Julie Kim and her laboratory for their expertise in organoid models and scaffold-free culture using agarose micromolds. Single cell library preparation and sequencing was done at Northwestern University NUSeq core facility with the support of NIH Grant (1S10OD025120). We would like to thank Dr. Jennifer Wai and Dr. Matthew Schipma for their assistance with single cell library preparation, sequencing, and bioinformatic analysis. Portions of Figures 1 and 3 were created using BioRender.com.

## Author contributions

S.S.D., M.Q.G., P.K., A.C., E.J.Z., J.M.P, and F.E.D. conceived the project and designed experiments. S.S.D., M.Q.G., P.K., A.C., E.J.Z., A.F., and T.C. performed experiments and analyzed data. S.S.D., M.Q.G., P.K., A.C., and E.J.Z. created figures and wrote the manuscript. All authors (S.S.D., M.Q.G., P.K., A.C., E.J.Z., A.F., T.C., M.T.P., and M.Z., J.M.P, and F.E.D.) edited and approved the final draft.

## Data availability

Single cell RNA-sequencing data files can be downloaded from NCBI Gene Expression Omnibus at accession number GSE273891.

---

*Supplemental Figure S1. Organoids remain intact when removed from agarose micromolds and can be generated from rhesus macaque ovarian tissue.* A) Representative images of ovarian somatic organoids at days 1, 3, and 5 of additional culture following removal from micromolds after 3 initial days of culture. Scale bars = 400 µm. B) Quantification of average organoid area over 5 additional days of culture following removal from micromolds. P<0.05 for a vs b and P<0.0001 for all other comparisons by one-way ANOVA. N=28 organoids. C) Representative scan of H&E-stained rhesus macaque ovarian tissue. Dashed line indicates separation of cortex (C) from medulla (M). Scale bar = 200 µm. D) Representative images of ovarian somatic organoids generated from rhesus macaque ovarian cortex (top) or medulla (bottom) at days 1, 3, and 5 of culture. Representative images of H&E-stained rhesus macaque organoid sections following 5 days in culture. Scale bars for transmitted light images = 400 µm and scale bars for H&E images = 100 µm. N=1 for ovarian somatic cell isolation and ovarian somatic organoid generation from rhesus macaque ovarian tissue.

*Supplemental Figure S2. Relative abundance of some key cell populations changes in ovarian somatic organoids over time in culture.* A-F) Representative images of vimentin (A), alpha-smooth muscle actin (11-SMA, B), desmin (C), F4/80 (D), FOXL2 (E), and 3β-HSD (F) IHC staining of mouse ovarian tissue sections and ovarian somatic organoid sections at days 1 and 6 of culture. Arrows show F4/80-positive macrophages restricted to the perimeter of organoids (black, D) and 3β-HSD-positive steroidogenic cells absent from the core of organoids (white, F). Scale bars = 20 µm. Insets are thresholded images that show positive staining. Quantification of the positive area for each marker in regions of interest in mouse ovaries (N=3 ovaries) and ovarian somatic organoids at days 1 (N=36-45) and 6 of culture (N=39-49). *P<0.05, **P<0.01, and ****P<0.0001 by Kruskal-Wallis tests followed by Dunn’s multiple comparisons tests.

*Supplemental Figure S3. Epithelial, Luteal 2, and Granulosa 1 clusters exhibit the greatest age-dependent changes in gene expression.* A) UMAP plots separated for young and old monolayer cells and cells dissociated from young and old ovarian somatic organoids at day 1 and day 6 of culture. B) Violin plots of a marker gene for each cell cluster for the UMAP plot of primary mouse ovarian somatic cells from reproductively young (6-12 weeks) and reproductively old (10-14 months) mice following plating in 2D, prior to organoid generation (monolayer) and cells dissociated from young and old ovarian somatic organoids at days 1 and 6 of culture combined. C) Table of percentage of each cell cluster for monolayer, day 1 organoids, and day 6 organoids for each age. Bottom row of table includes the total number of cells for monolayer, day 1 organoids, and day 6 organoids for each age. D) Quantification of the number of differentially expressed genes (DEGs) with age within each cluster for cells dissociated from ovarian somatic organoids at day 6 of culture. E) Gene ontology (GO) analysis of differentially expressed genes in Epithelial, Granulosa 1, and Luteal 2 clusters with age for cells dissociated from ovarian somatic organoids at day 6 of culture. Pathways upregulated (Up) and downregulated (Down) with age are labelled and displayed in manually grouped categories.

*Supplemental Figure S4. Cells in the Stroma/Mesenchyme 1 cluster are matrix fibroblasts.* A) Violin plots of a marker gene for each cell cluster for the UMAP plot of only primary mouse ovarian somatic cells from reproductively young (6-12 weeks) and reproductively old (10-14 months) mice following plating in 2D, prior to organoid generation (monolayer).

*Supplemental Figure S5. Enriched morphological clusters are variable between different groups of old mice.* A) Contour overlays on the morphological UMAP space showing enriched morphologies for primary mouse ovarian somatic cells from young (purple) and old (red) mice for 4 independent trials.

*Supplemental Figure S6. LatA treatment improves the age-dependent impairment in organoid aggregation in most replicates.* A-B) Representative images of organoids generated from young and old primary ovarian somatic cells following treatment with Latrunculin A (LatA) or vehicle control at 1 hour and 9 hours post seeding into agarose micromolds for 2 additional trials to that shown in Figure 6F. Scale bars = 500 µm. Right panels for each time point are thresholded masks of cells in agarose micromolds.

## References

1. Kinnear, H.M. et al. The ovarian stroma as a new frontier. Reproduction 160, R25–r39 (2020).

2. Tingen, C.M. et al. A macrophage and theca cell-enriched stromal cell population influences growth and survival of immature murine follicles in vitro. Reproduction 141, 809–820 (2011).

3. Shen, L., Liu, J., Luo, A. & Wang, S. The stromal microenvironment and ovarian aging: mechanisms and therapeutic opportunities. Journal of Ovarian Research 16, 237 (2023).

4. Furuya, M. Ovarian cancer stroma: pathophysiology and the roles in cancer development. Cancers (Basel) 4, 701–724 (2012).

5. Isola, J.V.V. et al. A single-cell atlas of the aging mouse ovary. Nature Aging 4, 145–162 (2024).

6. Briley, S.M. et al. Reproductive age-associated fibrosis in the stroma of the mammalian ovary. Reproduction 152, 245–260 (2016).

7. Zhang, Z., Schlamp, F., Huang, L., Clark, H. & Brayboy, L. Inflammaging is associated with shifted macrophage ontogeny and polarization in the aging mouse ovary. Reproduction 159, 325–337 (2020).

8. Foley, K.G., Pritchard, M.T. & Duncan, F.E. Macrophage-derived multinucleated giant cells: hallmarks of the aging ovary. Reproduction 161, V5–v9 (2021).

9. Ben Yaakov, T., Wasserman, T., Aknin, E. & Savir, Y. Single-cell analysis of the aged ovarian immune system reveals a shift towards adaptive immunity and attenuated cell function. eLife 12, e74915 (2023).

10. Landry, D.A. et al. Metformin prevents age-associated ovarian fibrosis by modulating the immune landscape in female mice. Science Advances 8, eabq1475 (2022).

11. Amargant, F. et al. Ovarian stiffness increases with age in the mammalian ovary and depends on collagen and hyaluronan matrices. Aging Cell, e13259 (2020).

12. Umehara, T. et al. Female reproductive life span is extended by targeted removal of fibrotic collagen from the mouse ovary. Sci Adv 8, eabn4564 (2022).

13. Dipali, S.S. et al. Proteomic quantification of native and ECM-enriched mouse ovaries reveals an age-dependent fibro-inflammatory signature. Aging (Albany NY) 15, 10821–10855 (2023).

14. Ouni, E. et al. Proteome-wide and matrisome-specific atlas of the human ovary computes fertility biomarker candidates and open the way for precision oncofertility. Matrix Biol 109, 91–120 (2022).

15. Simon, L.E., Kumar, T.R. & Duncan, F.E. In vitro ovarian follicle growth: a comprehensive analysis of key protocol variables†. Biol Reprod 103, 455–470 (2020).

16. Gamwell, L.F., Collins, O. & Vanderhyden, B.C. The mouse ovarian surface epithelium contains a population of LY6A (SCA-1) expressing progenitor cells that are regulated by ovulation-associated factors. Biol Reprod 87, 80 (2012).

17. Mara, J.N. et al. Ovulation and ovarian wound healing are impaired with advanced reproductive age. Aging (Albany NY) 12, 9686–9713 (2020).

18. Zink, K.E., Dean, M., Burdette, J.E. & Sanchez, L.M. Imaging Mass Spectrometry Reveals Crosstalk between the Fallopian Tube and the Ovary that Drives Primary Metastasis of Ovarian Cancer. ACS Cent Sci 4, 1360–1370 (2018).

19. Alzamil, L., Nikolakopoulou, K. & Turco, M.Y. Organoid systems to study the human female reproductive tract and pregnancy. Cell Death & Differentiation 28, 35–51 (2021).

20. Haider, S. & Beristain, A.G. Human organoid systems in modeling reproductive tissue development, function, and disease. Human Reproduction 38, 1449–1463 (2023).

21. Kwong, J. et al. Inflammatory cytokine tumor necrosis factor alpha confers precancerous phenotype in an organoid model of normal human ovarian surface epithelial cells. Neoplasia 11, 529–541 (2009).

22. Kopper, O. et al. An organoid platform for ovarian cancer captures intra- and interpatient heterogeneity. Nat Med 25, 838–849 (2019).

23. Pierson Smela, M.D., et al. Directed differentiation of human iPSCs to functional ovarian granulosa-like cells via transcription factor overexpression. eLife 12, e83291 (2023).

24. Parkes, W.S. et al. Hyaluronan and Collagen Are Prominent Extracellular Matrix Components in Bovine and Porcine Ovaries. Genes (Basel) 12 (2021).

25. Amargant, F., Vieira, C., Pritchard, M.T. & Duncan, F.E. Systemic low-dose anti-fibrotic treatment attenuates ovarian aging in the mouse. bioRxiv, 2024.2006.2021.600035 (2024).

26. Tsukui, T. et al. Collagen-producing lung cell atlas identifies multiple subsets with distinct localization and relevance to fibrosis. Nat Commun 11, 1920 (2020).

27. Bochaton-Piallat, M.L., Gabbiani, G. & Hinz, B. The myofibroblast in wound healing and fibrosis: answered and unanswered questions. F1000Res 5 (2016).

28. Ding, H., Chen, J., Qin, J., Chen, R. & Yi, Z. TGF-β-induced α-SMA expression is mediated by C/EBPβ acetylation in human alveolar epithelial cells. Molecular Medicine 27, 22 (2021).

29. Nelson, J.F., Felicio, L.S., Osterburg, H.H. & Finch, C.E. Altered Profiles of Estradiol and Progesterone Associated with Prolonged Estrous Cycles and Persistent Vaginal Cornification in Aging C578L/6J Mice1. Biology of Reproduction 24, 784–794 (1981).

30. Morris, M.E. et al. A single-cell atlas of the cycling murine ovary. Elife 11 (2022).

31. Ben Yaakov, T., Wasserman, T., Aknin, E. & Savir, Y. Single-cell analysis of the aged ovarian immune system reveals a shift towards adaptive immunity and attenuated cell function. Elife 12 (2023).

32. Hernandez-Gonzalez, I. et al. Gene expression profiles of cumulus cell oocyte complexes during ovulation reveal cumulus cells express neuronal and immune-related genes: does this expand their role in the ovulation process? Mol Endocrinol 20, 1300–1321 (2006).

33. Phillip, J.M., Aifuwa, I., Walston, J. & Wirtz, D. The Mechanobiology of Aging. Annu Rev Biomed Eng 17, 113–141 (2015).

34. Pollard, T.D. & Cooper, J.A. Actin, a central player in cell shape and movement. Science 326, 1208–1212 (2009).

35. Chalut, Kevin J. & Paluch, Ewa K. The Actin Cortex: A Bridge between Cell Shape and Function. Developmental Cell 38, 571–573 (2016).

36. Wu, P.H. et al. Single-cell morphology encodes metastatic potential. Sci Adv 6, eaaw6938 (2020).

37. Nguyen, A., Yoshida, M., Goodarzi, H. & Tavazoie, S.F. Highly variable cancer subpopulations that exhibit enhanced transcriptome variability and metastatic fitness. Nat Commun 7, 11246 (2016).

38. Yin, Z. et al. A screen for morphological complexity identifies regulators of switch-like transitions between discrete cell shapes. Nat Cell Biol 15, 860–871 (2013).

39. Phillip, J.M. et al. Biophysical and biomolecular determination of cellular age in humans. Nature Biomedical Engineering 1, 0093 (2017).

40. Wu, P.-H. et al. Evolution of cellular morpho-phenotypes in cancer metastasis. Scientific Reports 5, 18437 (2015).

41. Bajwa, P. et al. Age related increase in mTOR activity contributes to the pathological changes in ovarian surface epithelium. Oncotarget 7, 19214–19227 (2016).

42. Fujiwara, I., Zweifel, M.E., Courtemanche, N. & Pollard, T.D. Latrunculin A Accelerates Actin Filament Depolymerization in Addition to Sequestering Actin Monomers. Curr Biol 28, 3183–3192.e3182 (2018).

43. Broekmans, F.J., Soules, M.R. & Fauser, B.C. Ovarian aging: mechanisms and clinical consequences. Endocr Rev 30, 465–493 (2009).

44. Auersperg, N., Wong, A.S.T., Choi, K.-C., Kang, S.K. & Leung, P.C.K. Ovarian Surface Epithelium: Biology, Endocrinology, and Pathology*. Endocrine Reviews 22, 255–288 (2001).

45. Thung, P.J., Boot, L.M. & Mühlbock, O. Senile Changes in the Oestrous Cycle and in Ovarian Structure in Some Inbred Strains of Mice. Acta Endocrinologica 23, 8–32 (1956).

46. Faddy, M.J., Jones, E.C. & Edwards, R.G. An analytical model for ovarian follicle dynamics. J Exp Zool 197, 173–185 (1976).

47. Lliberos, C. et al. Evaluation of inflammation and follicle depletion during ovarian ageing in mice. Sci Rep 11, 278 (2021).

48. Klingberg, F., Hinz, B. & White, E.S. The myofibroblast matrix: implications for tissue repair and fibrosis. J Pathol 229, 298–309 (2013).

49. Mederacke, I., Dapito, D.H., Affò, S., Uchinami, H. & Schwabe, R.F. High-yield and high-purity isolation of hepatic stellate cells from normal and fibrotic mouse livers. Nature Protocols 10, 305–315 (2015).

50. Berdyyeva, T.K., Woodworth, C.D. & Sokolov, I. Human epithelial cells increase their rigidity with ageing in vitro: direct measurements. Phys Med Biol 50, 81–92 (2005).

51. Dulińska-Molak, I. et al. Age-Related Changes in the Mechanical Properties of Human Fibroblasts and Its Prospective Reversal After Anti-Wrinkle Tripeptide Treatment. Int J Pept Res Ther 20, 77–85 (2014).

52. Schulze, C. et al. Stiffening of human skin fibroblasts with age. Clin Plast Surg 39, 9–20 (2012).

53. Esue, O., Tseng, Y. & Wirtz, D. Alpha-actinin and filamin cooperatively enhance the stiffness of actin filament networks. PLoS One 4, e4411 (2009).

54. Starodubtseva, M.N. Mechanical properties of cells and ageing. Ageing Res Rev 10, 16–25 (2011).

55. Xu, W. et al. Cell stiffness is a biomarker of the metastatic potential of ovarian cancer cells. PLoS One 7, e46609 (2012).

56. Rowley, J.E. et al. Low Molecular Weight Hyaluronan Induces an Inflammatory Response in Ovarian Stromal Cells and Impairs Gamete Development In Vitro. Int J Mol Sci 21 (2020).

57. Rowley, J.E. et al. Tissue-specific Fixation Methods Are Required for Optimal In Situ Visualization of Hyaluronan in the Ovary, Kidney, and Liver. J Histochem Cytochem 68, 75–91 (2020).

58. Hao, Y. et al. Integrated analysis of multimodal single-cell data. Cell 184, 3573–3587.e3529 (2021).

59. Chen, E.Y. et al. Enrichr: interactive and collaborative HTML5 gene list enrichment analysis tool. BMC Bioinformatics 14, 128 (2013).

60. Spandidos, A., Wang, X., Wang, H. & Seed, B. PrimerBank: a resource of human and mouse PCR primer pairs for gene expression detection and quantification. Nucleic Acids Res 38, D792–799 (2010).

61. Livak, K.J. & Schmittgen, T.D. Analysis of relative gene expression data using real-time quantitative PCR and the 2(-Delta Delta C(T)) Method. Methods 25, 402–408 (2001).

62. Sayers, E.W. et al. Database resources of the national center for biotechnology information. Nucleic Acids Res 50, D20–d26 (2022).

