## Supplemental Figures for "Self-organizing ovarian somatic organoids preserve cellular heterogeneity and reveal cellular contributions to ovarian aging"

### Supplemental Figure S1

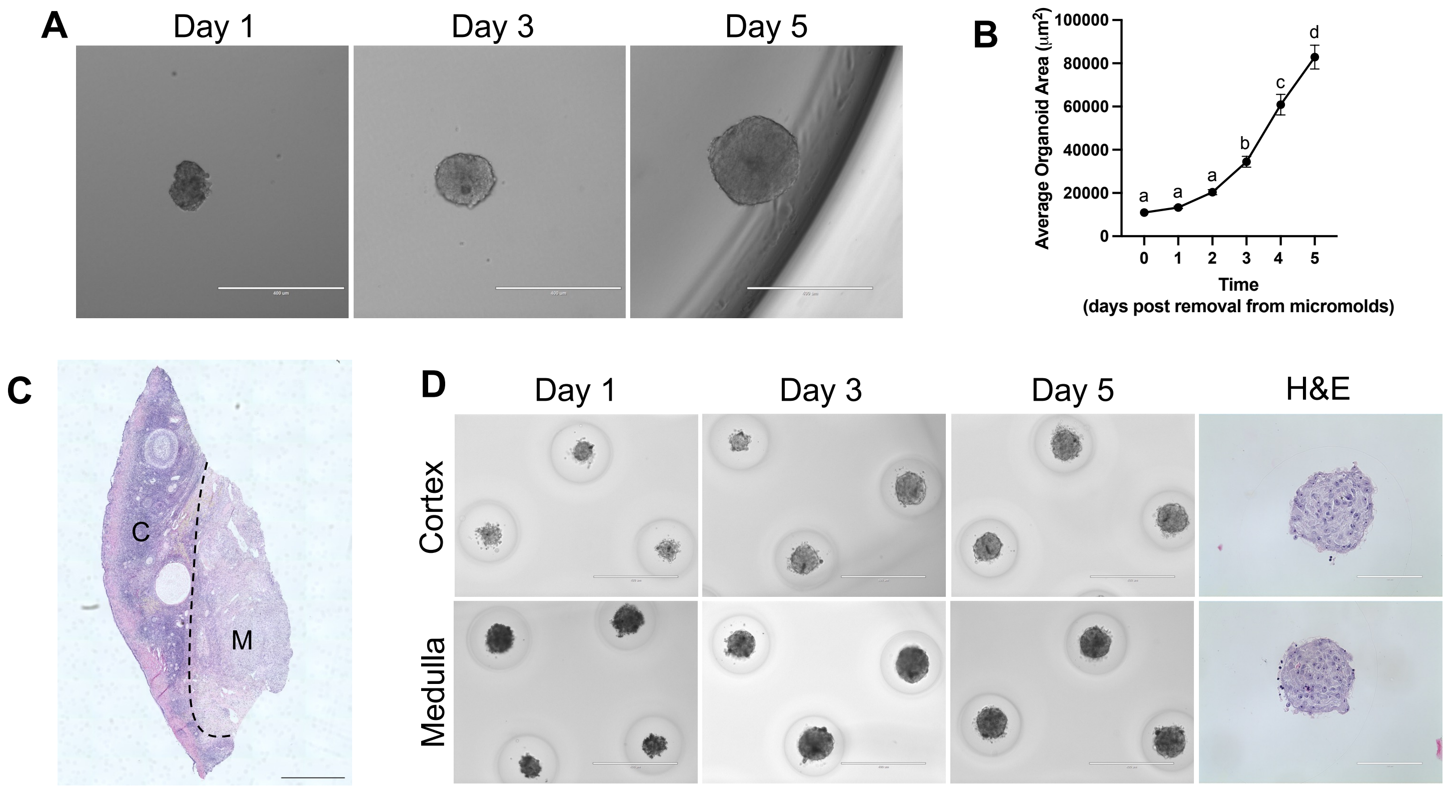

### Supplemental Figure S2

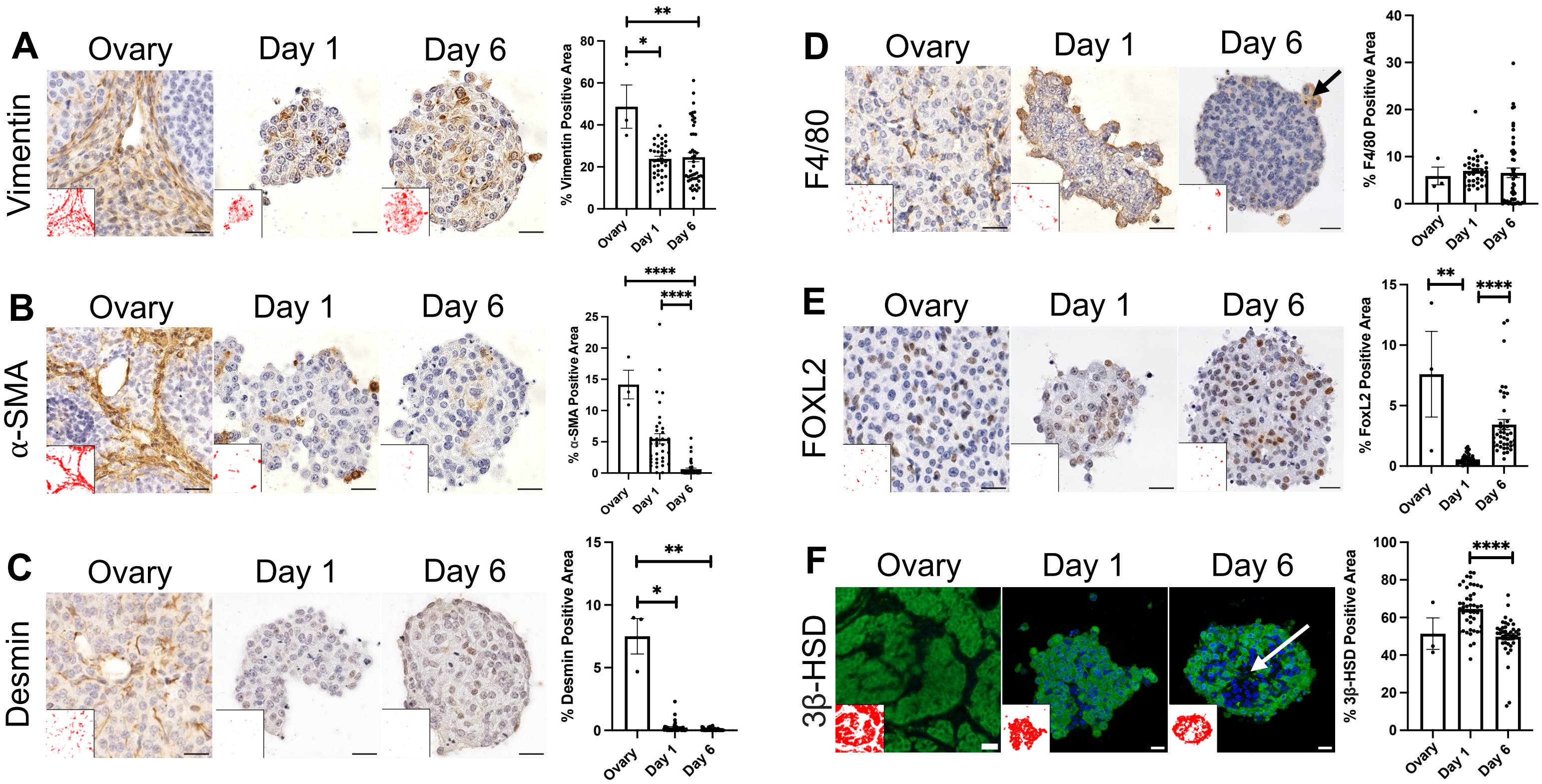

### Supplemental Figure S3

A

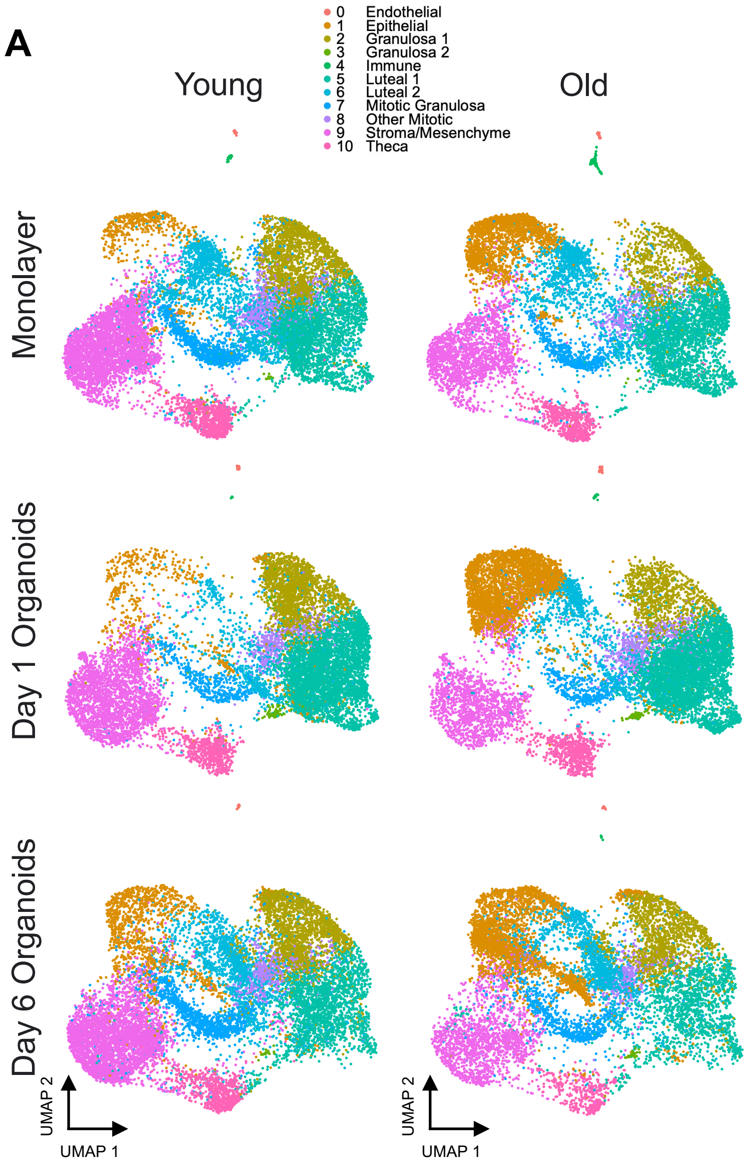


B

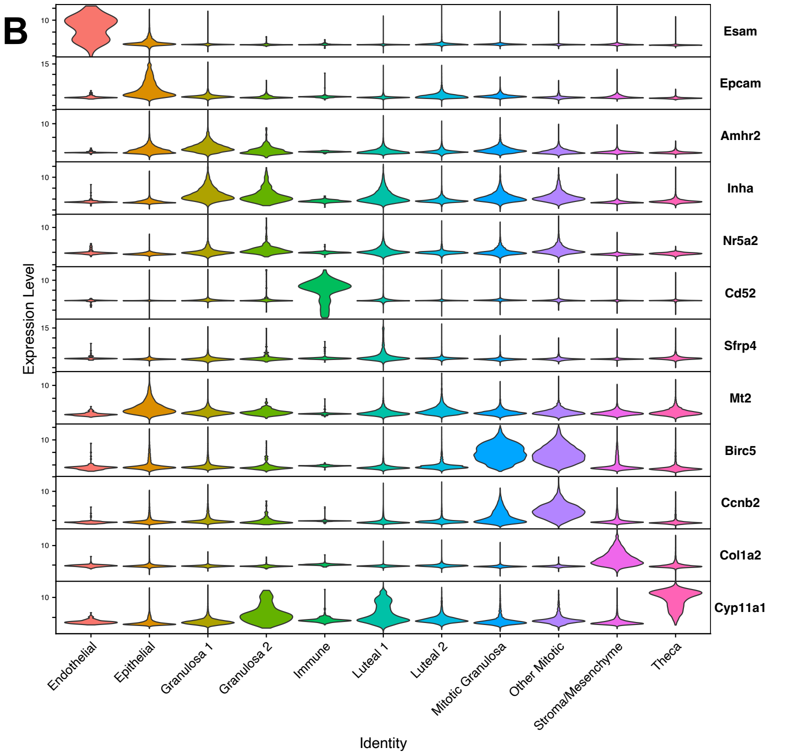


C

| Cluster | Young Monolayer | Young D1 | Young D6 | Old Monolayer | Old D1 | Old D6 |
|---|---|---|---|---|---|---|
| 0 | 0.1 ± 0.0% | 0.2 ± 0.2% | 0.1 ± 0.0% | 0.3 ± 0.0% | 0.3 ± 0.0% | 0.1 ± 0.0% |
| 1 | 4.2 ± 1.1% | 4.6 ± 0.1% | 8.9 ± 1.5% | 19.5 ± 3.0% | 25.1 ± 5.2% | 28.9 ± 1.0% |
| 2 | 14.2 ± 0.1% | 15.9 ± 3.2% | 14.1 ± 3.2% | 8.5 ± 0.2% | 7.5 ± 1.2% | 12.9 ± 1.0% |
| 3 | 0.1 ± 0.0% | 0.8 ± 0.1% | 0.1 ± 0.1% | 0.0 ± 0.0% | 0.5 ± 0.1% | 0.4 ± 0.2% |
| 4 | 0.2 ± 0.1% | 0.0 ± 0.0% | 0.0 ± 0.0% | 1.1 ± 0.4% | 0.2 ± 0.0% | 0.1 ± 0.0% |
| 5 | 20.9 ± 1.4% | 37.2 ± 3.5% | 17.1 ± 5.2% | 24.7 ± 1.3% | 32.9 ± 2.6% | 14.4 ± 2.2% |
| 6 | 14.5 ± 0.2% | 6.3 ± 0.8% | 12.3 ± 2.5% | 14.8 ± 2.0% | 9.7 ± 2.0% | 16.0 ± 1.7% |
| 7 | 8.0 ± 1.8% | 4.1 ± 0.2% | 7.9 ± 2.7% | 5.7 ± 1.2% | 3.0 ± 0.3% | 6.3 ± 1.3% |
| 8 | 4.9 ± 0.7% | 4.0 ± 0.0% | 4.0 ± 1.2% | 2.5 ± 0.9% | 3.9 ± 0.6% | 3.8 ± 0.0% |
| 9 | 25.9 ± 3.6% | 21.5 ± 4.9% | 29.8 ± 4.4% | 16.4 ± 1.5% | 11.8 ± 1.7% | 13.7 ± 2.4% |
| 10 | 6.9 ± 1.1% | 5.4 ± 1.0% | 5.8 ± 1.6% | 6.4 ± 0.1% | 5.2 ± 0.1% | 3.3 ± 0.4% |
| Total Number of Cells | 13873 | 12297 | 13477 | 10994 | 13470 | 10964 |

D

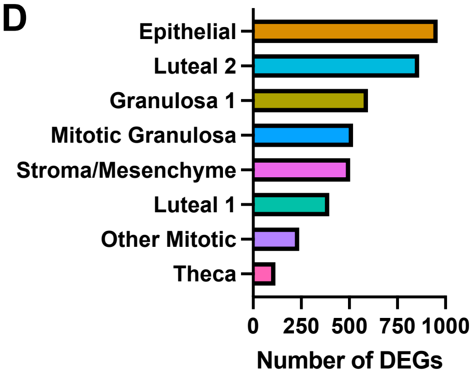


E

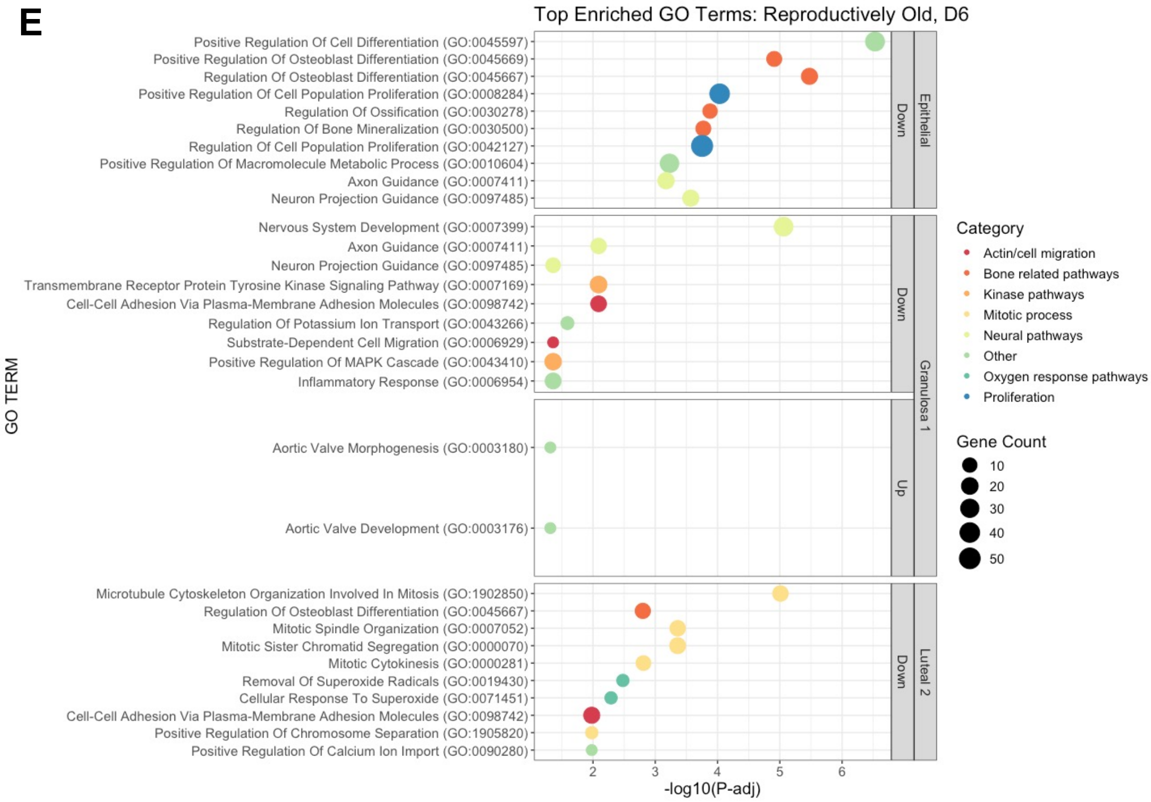

### Supplemental Figure S4

**A**

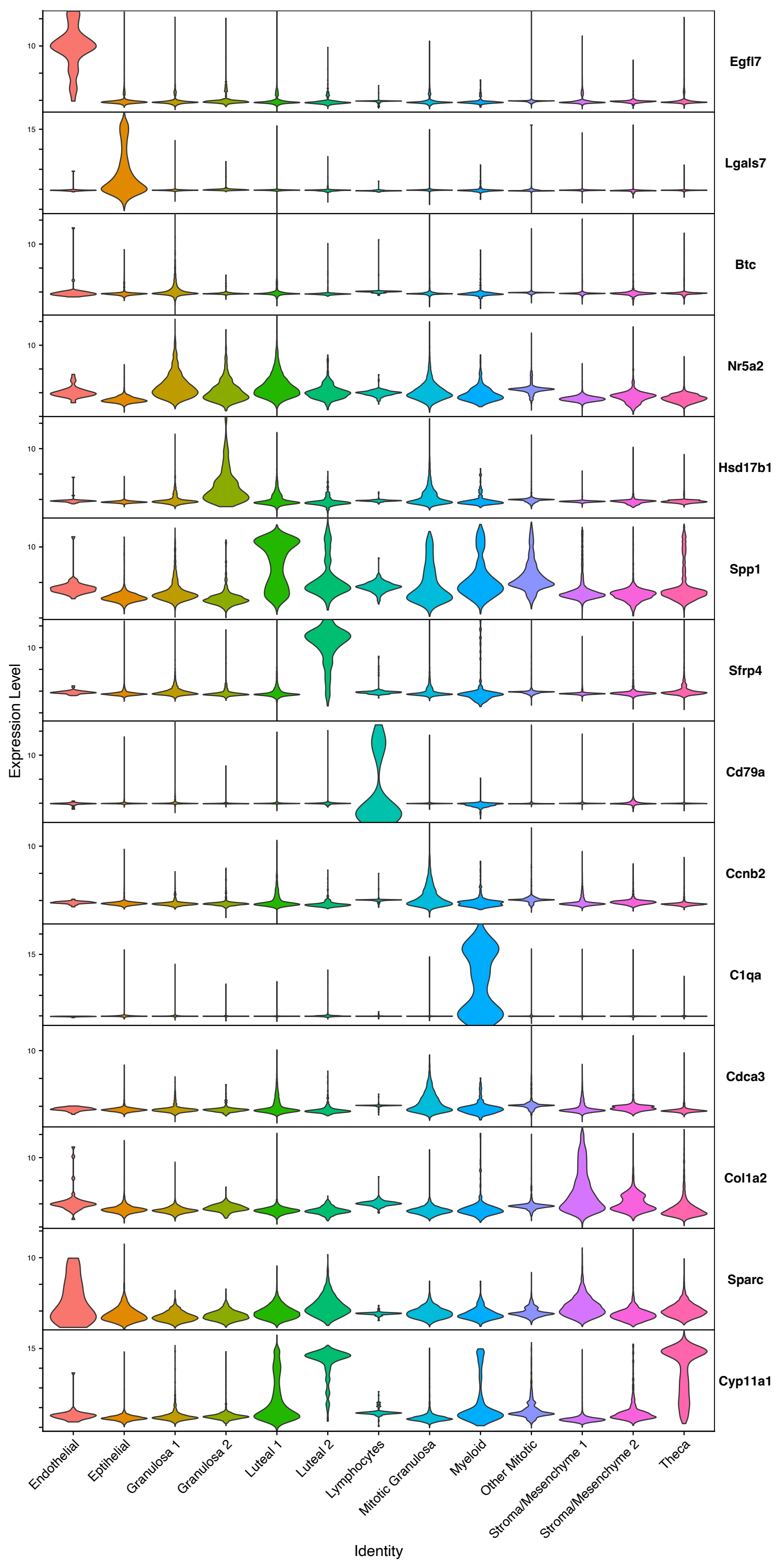

### Supplemental Figure S5

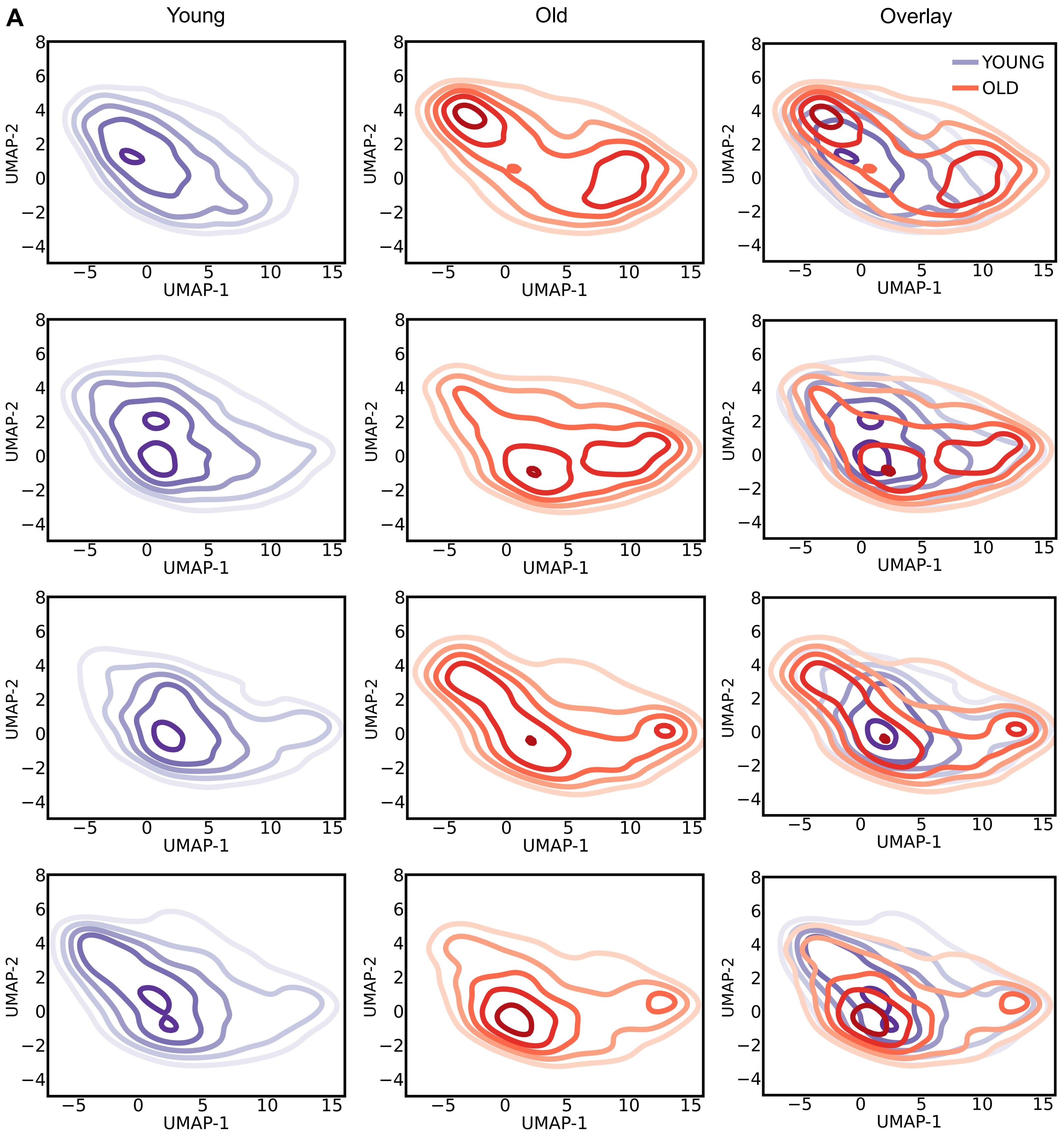

### Supplemental Figure S6

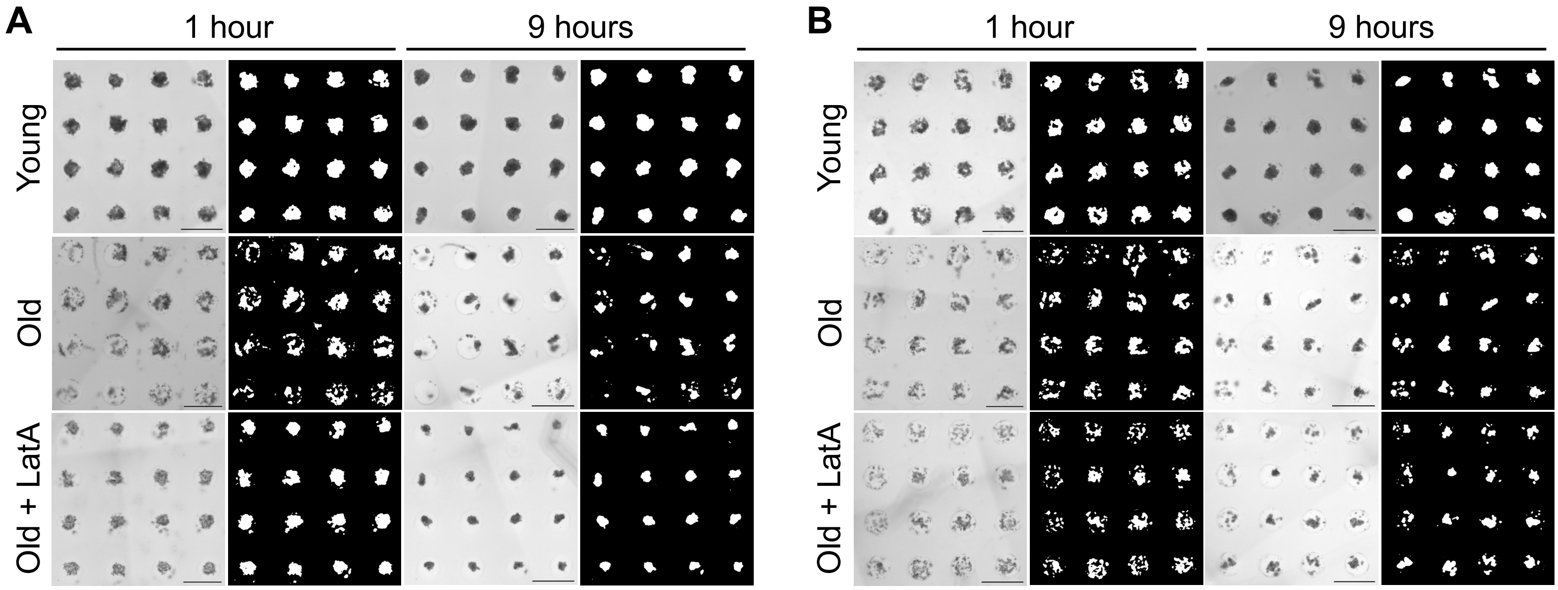
