## Supplemental Tables for "Self-organizing ovarian somatic organoids preserve cellular heterogeneity and reveal cellular contributions to ovarian aging"

**Supplemental Table S1. Statistically significant changes in cellular composition of organoids generated from reproductively young and old mice over six days in culture.**

| **Marker** | **Comparison** | **P-value** |
| --- | --- | --- |
| Vimentin | Young Day 1 vs Old Day 6 | P<0.0001 |
| ⍺-SMA | Old Day 1 vs Young Day 6 | P<0.0001 |
| F4/80 | Old Day 1 vs Young Day 6 | P=0.0068 |
| FOXL2 | Old Day 1 vs Young Day 6 | P<0.0001 |
| 3β-HSD | Young Day 1 vs Old Day 6 | P<0.0001 |
| 3β-HSD | Old Day 1 vs Young Day 6 | P<0.0001 |

**Supplemental Table S2. Antibodies used for Immunolabeling.**

| **Antibody** | **Host Species** | **Concentration** | **Manufacturer/**  **Catalog Number** |
| --- | --- | --- | --- |
| **Primary Antibodies** | | | |
| Ki67 | Rabbit | 1:400 | Cell Signaling Technology / 12202 |
| CC3 | Rabbit | 1:250 | Cell Signaling Technology / 9579T |
| Vimentin | Rabbit | 1:100 (IHC)  1:50 (ICC) | Cell Signaling Technology / 5741S |
| ⍺-SMA | Rabbit | 1:500 | Cell Signaling Technology / 19245 |
| 3β-HSD | Rabbit | 1:500 | Cosmo Bio / K0607 |
| FOXL2 | Rabbit | 1:200 | Abcam / ab246511 |
| F4/80 | Rat | 1:50 | Bio-Rad / MCA497G |
| **Secondary Antibodies** | | | |
| Biotinylated Anti-Rabbit IgG | Goat | 1:200 (IHC) | Vector Laboratories / PK-6101 |
| Biotinylated Anti-Rat IgG | Goat | 1:100 (IHC) | Vector Laboratories / BA-9400-1.5 |
| Anti-Rabbit Alexa Fluor 488 | Goat | 1:100 (ICC) | Thermo Scientific / A-11004 |

**Supplemental Table S3. RT-qPCR Primer Sequences (5’-3’).**

| **Primer** | **Forward** | **Reverse** |
| --- | --- | --- |
| *Acta2* | GTCCCAGACATCAGGGAGTAA | TCGGATACTTCAGCGTCAGGA |
| *Col1a1* | ATGTTCAGCTTTGTGGACCTC | CAGAAAGCACAGCACTCGC |
| *Gapdh* | GCACAGTCAAGGCCGAGAAT | GCCTTCTCCATGGTGGTGAA |
